# Cooperation between CYB5R3 and NOX4 via coenzyme Q mitigates endothelial inflammation

**DOI:** 10.1101/2021.08.12.456058

**Authors:** Shuai Yuan, Scott A. Hahn, Megan P. Miller, Subramaniam Sanker, Michael J Calderon, Mara Sullivan, Atinuke M. Dosunmu-Ogunbi, Marco Fazzari, Yao Li, Michael Reynolds, Katherine C Wood, Claudette M. St. Croix, Donna Stolz, Eugenia Cifuentes-Pagano, Placido Navas, Sruti Shiva, Francisco J. Schopfer, Patrick J. Pagano, Adam C. Straub

**Affiliations:** Heart, Lung, Blood and Vascular Medicine Institute, University of Pittsburgh, Pittsburgh, Pennsylvania; Department of Pharmacology and Chemical Biology, University of Pittsburgh, Pittsburgh, Pennsylvania; Center for Biologic Imaging, University of Pittsburgh, Pittsburgh, Pennsylvania; Centro Andaluz de Biología del Desarrollo and CIBERER, Instituto de Salud Carlos III, Universidad Pablo de Olavide-CSIC-JA, Sevilla, Spain, Spain; Pittsburgh Liver Research Center (PLRC), University of Pittsburgh, Pittsburgh, Pennsylvania; Center for Microvascular Research, University of Pittsburgh, Pittsburgh, Pennsylvania

**Keywords:** CYB5R3, NOX4, CoQ, ROS, inflammation

## Abstract

NADPH oxidase 4 (NOX4) regulates endothelial inflammation by producing reactive oxygen species. Since coenzyme Q (CoQ) mimics affect NOX4 activity, we hypothesize that cytochrome b5 reductase 3 (CYB5R3), a CoQ reductase abundant in vascular endothelial cells, modulates inflammatory activation.

Mice lacking endothelial CYB5R3 (R3 KO), under lipopolysaccharides (LPS) challenge, showed exacerbated hypotension, decreased acetylcholine-induced vasodilation, and elevated vascular adhesion molecule 1 (Vcam-1) mRNA in aorta. In vitro, silencing *Cyb5r3* enhanced LPS-induced VCAM-1 protein in a NOX4 dependent manner. APEX2- based electron microscopy and proximity biotinylation demonstrated CYB5R3’s localization on the mitochondrial outer membrane and its interaction with NOX4, which was further confirmed by the proximity ligation assay. Notably, *Cyb5r3* silenced HAECs had less total H_2_O_2_ but more mitochondrial O_2_^•-^. Using inactive or non-membrane bound active CYB5R3, we found CYB5R3 activity and membrane translocation were needed for optimal generation of H_2_O_2_ by NOX4. Lastly, CoQ deficient cells showed decreased NOX4-derived H_2_O_2_, indicating a requirement for endogenous CoQ in NOX4 activity.

In conclusion, CYB5R3 mitigates endothelial inflammatory activation by assisting in NOX4-dependent H_2_O_2_ generation via CoQ.

**NOVELTY AND SIGNIFICANCE:** *What Is Known?:* NADPH oxidase 4 (NOX4) reportedly produces primarily hydrogen peroxide (H_2_O_2_) and, to a lesser extent, superoxide (O_2_^•-^) and has been shown to have both beneficial and deleterious effects in the cardiovascular system. NOX4 activity can be affected by NAD(P)H quinone oxidoreductase 1 (NQO1), a CoQ reductase, and synthetic quinone compounds used to mimic CoQ. Cytochrome b5 reductase 3 (CYB5R3) is known to reduce CoQ and is highly expressed in endothelial cells.

*What New Information Does This Article Contribute?:* In vivo, the lack of endothelial CYB5R3 causes exacerbated lipopolysaccharides (LPS)-induced inflammatory signaling, endothelial dysfunction, and hypotension. Endothelial CYB5R3 mitigates inflammatory signaling by LPS and tumor necrosis factor α (TNF-α) in a NOX4 dependent manner. In endothelial cells, CYB5R3 and NOX4 reside in close proximity on the mitochondrial outer membrane. NOX4’s ability to generate H_2_O_2_ depends on the membrane translocation and activity of CYB5R3 and the presence of endogenous CoQ.

*NONSTANDARD Abbreviations and Acronyms:* 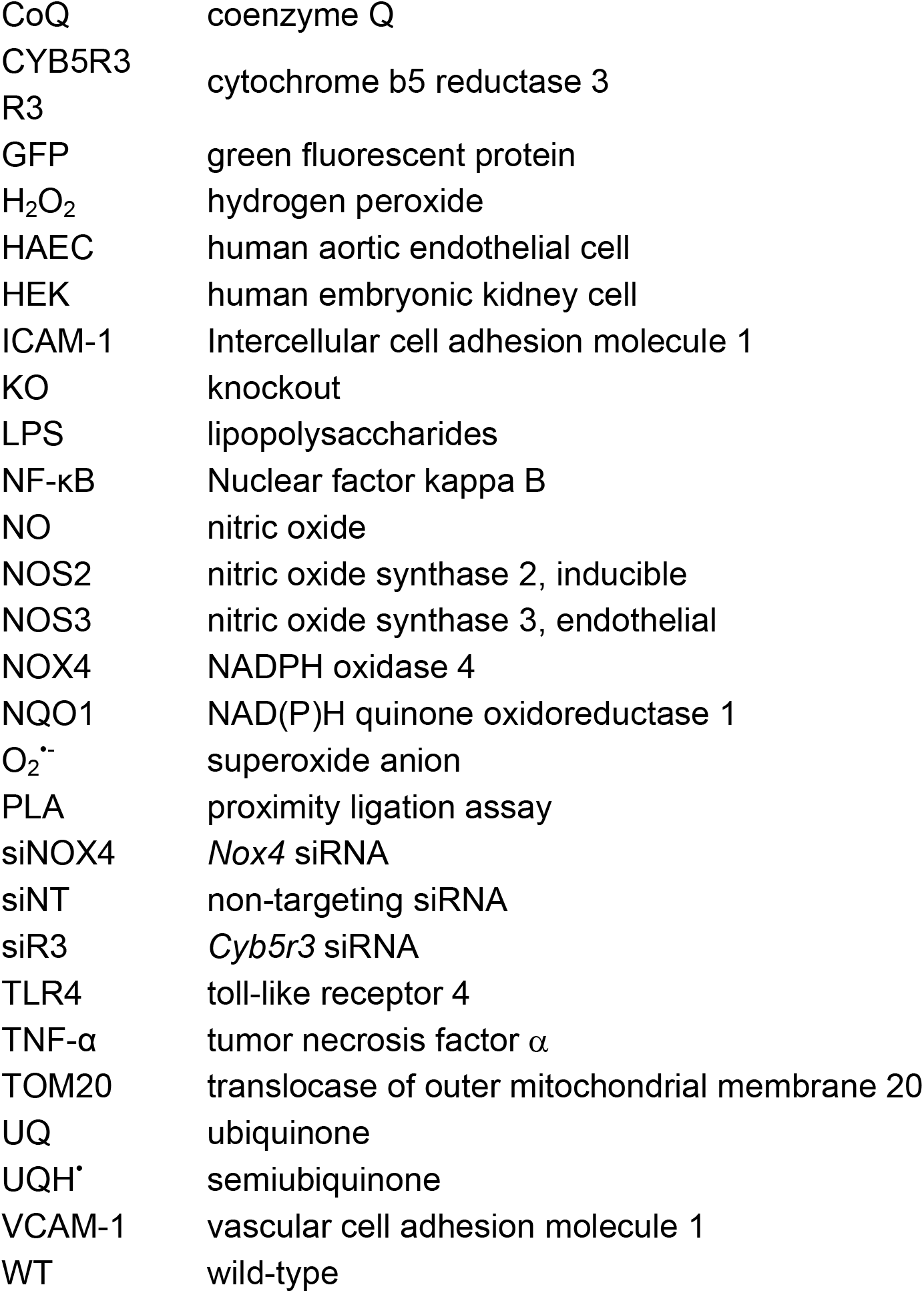 Protein names are abbreviated as capital letters (e.g., CYB5R3), while the corresponding gene names are annotated as in italic lower cases (e.g., *Cyb5r3*).

## INTRODUCTION

In the vascular wall, the endothelium governs leukocyte recruitment during inflammation to help detain and remediate pathogens^1^. Pathogenic stimuli such as endotoxin activate endothelium to rapidly deploy adhesion molecules to the surface to entrap leukocytes and facilitate diapedesis^2^. Vascular cell and intercellular adhesion molecules, VCAM-1 and ICAM-1, are major endothelial recruitment factors for leukocytes at sites of infection^3–5^. In severe infection, endotoxin released by circulating pathogens exacerbates this process, causing systemic hypotension, reduced peripheral blood flow, and multi-organ failure^6, 7^. One important element of the inflammatory signaling response is reactive oxygen species. Reactive oxygen species are being increasingly appreciated for their essential role in cellular signal transduction and conditioning. On the one hand, phagosomal superoxide (O_2_^•-^) in leukocytes kills bacteria, whereas excessive reactive species such as O_2_^•-^, hydrogen peroxide (H_2_O_2_), and peroxynitrite can contribute to host endothelial overactivation and dysfunction^8, 9^.

Endothelial cells employ a diversity of machinery, under physiological and pathological conditions, to generate reactive species such as O_2_^•-^ and H_2_O_2_. NADPH oxidases (NOX) are a family of enzymes dedicated to reducing ambient oxygen for O_2_^•-^ and H_2_O_2_ formation. In the NOX family, NOX4 is unique on several levels: 1) it is constitutively activated, 2) it predominantly produces H_2_O_2_ versus O_2_^•-^, 3) it is compartmentalized on intracellular membranes and in particular the mitochondrion and 4) it serves divergent roles in the cardiovascular system^10–12^. Although NOX4 is protective in animal models of chronic inflammation, including atherosclerosis and peripheral arterial disease^13, 14^, it exacerbates the acute inflammatory response induced by lipopolysaccharides and tumor necrosis factor α (TNF-α) in endothelial cells^15–17^. However, it remains incompletely understood how NOX4 causes these opposing cardiovascular outcomes.

One explanation for NOX4’s dichotomous functions in cardiovascular outcomes likely stems from its capacity to generate two different primary products, H_2_O_2_ and O_2_^•-^. Coenzyme Q (CoQ) has been suggested to augment NOX4’s production of H_2_O_2,_ but the evidence is limited to biochemical studies using synthetic quinone compounds and a CoQ reductase, NAD(P)H quinone oxidoreductase 1 (NQO1)^18^. Ubiquitously synthesized by mammalian cells, CoQ has a long hydrophobic side chain embedded in the lipid bilayer and a benzoquinone head group. The head group is redox-sensitive and can receive or donate up to two electrons to cycle between three distinct states: the fully oxidized ubiquinone, the fully reduced ubiquinol, and the semiubiquinone radical. CoQ redox cycling allows it to shuttle electrons between enzymes maintaining their activity^19^. Therefore, endogenous CoQ may modulate NOX4 activity with the assistance of a CoQ reductase.

Cytochrome b5 reductase 3 (CYB5R3) is a CoQ reductase highly expressed in endothelial cells. CYB5R3-dependent CoQ reduction is essential in the membrane antioxidation pathway^20–22^. In addition to CoQ reduction, CYB5R3 mediates the reduction of hemeproteins such as cytochrome b5 and hemoglobin, which is required for fatty acid metabolism and the prevention of methemoglobinemia^23, 24^. We have previously demonstrated that endothelial cells in small arteries and arterioles use CYB5R3 to modulate nitric oxide signaling via α-globin heme redox cycling^25^. However, the role of endothelial cell CYB5R3 in large arteries, such as the aorta, has not been investigated. This constitutes a critical research gap since endothelial cells of large arteries abundantly express CYB5R3 but do not express α-globin or use fatty acid as a primary source of energy. Therefore, considering the critical role of NOX4 in inflammation and the potential interaction between NOX4 and CYB5R3, we hypothesized that endothelial CYB5R3 regulates inflammatory activation by modulating NOX4 oxidant formation.

## METHODS AND MATERIALS

### Animals

All animal studies were approved by and in compliance with the University of Pittsburgh Institutional Animal Care and Use Committee. *Cyb5r3* floxed mice (*Cyb5r3^fl/fl^)* were generated as previously described^26^. Both wild-type C57BL/6J and *Cyb5r3^fl/fl^* mice were crossed with the tamoxifen-inducible *Cdh5(PAC)-CreERT2* mice kindly provided by Dr. Ralf Adams^27^. Male mice at the age of 10-12 weeks were treated with tamoxifen (10 mg/kg in corn oil) intraperitoneally for five consecutive days and allowed to rest for seven days before experiments. All mice were assigned a number randomly at birth, and the genotype was concealed from the technician during LPS treatment and data acquiring. All mice used in this study were reported.

### Cell culture

Human aortic endothelial cells (HAEC) were purchased from Lonza and maintained in endothelial growth medium (EGM-2, Lonza, CC-3162), with 5% CO_2_, at 37°C. According to Lonza’s suggestion, population doubling was used to track the age of cells in culture instead of the conventional passage number. Cells less than 13 times of population doublings were used for experiments. In an experiment, HAECs were allowed to grow fully confluent before they were challenged with LPS (1 or 10 ug/ml) for indicated incubation time.

HEK293FT cells were obtained from Thermo Fisher Scientific (R70007) and grew in Dulbecco’s Modified Eagle Medium (DMEM, 12430062) with 10% fetal bovine serum. *COQ6* gene deletion was achieved in HEK293 cells with the CRISPR-Cas9 technology and kindly offered by Dr. Placido Navas^28^. The *COQ6* knockout cell line and corresponding control cell line were maintained in DMEM with 10% fetal bovine serum. Additionally, 10 µM uridine was supplied for both cell lines to compensate for the loss of CoQ.

### Cloning of expression vectors

The wild-type (WT) human *Cyb5r3* (NM_000398.6) was cloned from a pINCY vector (LIFESEQ1901142, Open Biosystems) into the pcDNA3.1 expression vector (Invitrogen, V86020). The nucleotides encoding the first 23 amino acids for membrane anchoring were deleted (ΔAA1-23) by PCR. Two inactive variants, K111A and K111A/G180V, were created by point mutations on the pcDNA3.1-*Cyb5r3 (WT)* plasmid using QuikChange II XL Site-Directed Mutagenesis Kit (Agilent, 200521). Mutagenesis primers were designed using Agilent’s online program. Additionally, human *Nox4* (AF254621.1) and *p22* (NM_000101.4) in pcDNA3.1 were kindly provided by Dr. Patrick Pagano.

### APEX2 related electron microscopy and affinity pulldown

Transient expression of CYB5R3-APEX2 was achieved by using lentivirus as described above. Our bicistronic vector coexpresses CYB5R3-APEX2 and green fluorescent protein (GFP) with an internal ribosome entry site (IRES). To avoid arbitrary effects due to a high level of expression, we used a low concentration of virus to obtain 10% GFP positive cells. APEX2 based electron microscopy and affinity enrichment were performed according to previous publications^29, 30^.

For electron microscopy, HAECs were fixed with 2% (v/v) glutaraldehyde in 100 mM cacodylate solution. Fixed cells were loaded with 0.5 mg/ml 3,3′-Diaminobenzidine (DAB) and treated with 10 mM H_2_O_2_. Once brown precipitates form in CYB5R3-APEX2 positive cells, cells were rinsed with 100 mM cacodylate solution, post-fixed in 1% osmium tetroxide with 1% potassium ferricyanide, rinsed in PBS, dehydrated through a graded series of ethanol, and embedded in Poly/Bed® 812 (Luft formulations). Semi- thin (300 nm) sections were cut on a Leica Reichart Ultracut, stained with 0.5% Toluidine Blue in 1% sodium borate, and examined under the light microscope. Ultrathin sections (65 nm) were examined on a JEOL 1400 Plus transmission electron microscope with a side mount AMT 2k digital camera (Advanced Microscopy Techniques, Danvers, MA).

For affinity pulldown, HAECs were incubated with 500 µM biotin-phenoxyl radical (Cayman, 26997) for 30 minutes. To activate APEX2, H_2_O_2_ was added at the final concentration of 100 µM for 60 seconds. Biotin-phenoxyl radical were immediately neutralized by repeated washing with the quench buffer (10 mM sodium ascorbate, 10 mM sodium azide, 5 mM Trolox [Cayman, 10011659] in phosphate-buffered saline, pH 8). Biotinylated proteins were enriched using Dynabeads™ M-280 Streptavidin (Invitrogen, 11205D) and eluted by heating the beads in 2X Laemmli buffer for immunoblotting.

### Quantification of H_2_O_2_ and NOX4 activity

Cellular production of H_2_O_2_ was evaluated by its fluorescent reaction product with coumarin boronate (Cayman, 14051) in the presence of L-NAME (Cayman, 80210) and taurine (Cayman, 27031)^31–33^. For endothelial cells, cells were plated and transfected in a 96-well plate. To measure H_2_O_2_, the medium was changed to the assay buffer (20 mM HEPES pH 7.5, 10 µM EDTA, 100 µM L-NAME, 1 mM taurine, and 0.01% bovine serum albumin in DMEM). For each group, negative controls were included by adding catalase to the final concentration of 500 µg/ml. Reactions were started by adding 20 µM CBA probe. Fluorescence (350/450 nm) was measured kinetically at 37 °C for four hours. At the end of the assay, endothelial cells were stained with crystal violet to determine the number of cells. To calculate the reaction rate, the log phase slope was normalized with the crystal violet staining results. For HEK293(FT) cells, since they did not withstand repeated rinsing during crystal violet staining, a different strategy was used for normalization. After transfection, HEK293(FT) cells were dislodged, counted, and resuspended in the assay buffer so that each well in a 96-well plate contains 120,000 cells at the time of the assay. Otherwise, the CBA assay was performed the same as for HAECs. Additionally, cells were pelleted from the remaining cell suspension for protein quantification to normalize the result. Importantly, for each treatment, the fluorescent signal from a corresponding catalase control was included, and the catalase-inhibitable amount was designated as H_2_O_2_.

### Mitochondrial O_2_^•-^ measurement

Cells were transfected in 10-cm dishes for siRNA transfection. To measure mitochondrial O_2_^•-^, cells were trypsinized and resuspended in Ca^2+^ and Mg^2+^ negative Hank’s balanced salt solution (HBSS). In a 96-well plate, 100,000 cells were added to each well with 5 µM MitoSOX™ Red (Invitrogen, M36008). Immediately after the addition of MitoSOX, the fluorescent signal (510/580 nm) was recorded at 37 °C for two hours. To reduce variation, fluorescent intensities within every five minutes were averaged before the log phase slope was calculated to represent the reaction rate. The results were normalized with the protein concentrations measured in the remaining cell suspension.

### Statistics

Data were reported as mean ± standard error of the mean unless specified otherwise. The n value represents the number of animals or independent experiments repeated on different days. Statistical analysis was performed with Graph Pad Prism 8 using Student t-test, one-way ANOVA, and two-way ANOVA with Sidak or Tukey post-hoc test. P-values less than 0.05 were considered as statistically significant. Specific tests used for experiments are shown in figure legends.

**Extended methods can be found in the supplementary material.**

## RESULTS

### Endothelial CYB5R3 protects against LPS-induced inflammatory responses *in vivo*

*Cyb5r3* floxed mice were previously generated by our group and crossed with *Cdh5- CreER^T^*^2^ mice to create the tamoxifen-inducible, endothelium-specific *Cyb5r3* knockout mice (R3 KO)^26^ (**Figure 1a**). After five consecutive days of intraperitoneal tamoxifen injections (10 mg/kg), endothelial CYB5R3 protein was absent in the aorta from R3 KO mice (**Figure 1a**). To determine whether endothelial CYB5R3 affects LPS induced systemic hypotension, we implanted radiotelemetry units in mice to monitor real-time changes in arterial blood pressure. At baseline, there was no difference in resting blood systolic pressure between R3 KO mice and WT controls during a full 24-hour cycle (**Figure 1b**). However, significantly lower heart rates were observed in the R3 KO mice (**Supplemental Table 1**). A single sublethal dose of LPS (5 mg/kg) injected intraperitoneally caused a drop in systolic blood pressure in WT (**Figure 1c**). Importantly, R3 KO mice suffered a more significant loss of systolic blood pressure (34.4±17 vs. 20.2±19.5 mmHg at the 6-hour time point). In a separate cohort of mice, aortae were collected from WT and R3 KO mice 8 hours after LPS (5 mg/kg) injections. *Ex vivo* wire myography showed inhibited acetylcholine-induced vasodilation in R3 KO aortic rings, suggesting LPS resulted in more severe endothelial dysfunction in the *Cyb5r3* deficient endothelium (**Figure 1d**). It is worth noting that the vasodilation induced by acetylcholine *ex vivo* depends on endothelial nitric oxide synthase (NOS3) function^34^. In contrast, LPS-induced vasodilation *in vivo* is known to be mediated by nitric oxide derived from inducible nitric oxide synthase (NOS2) in leukocytes^35, 36^. Indeed, inflammatory genes such as *Nos2* and vascular cell adhesion molecule 1 (*Vcam1*), but not intercellular adhesion molecule 1 (*Icam1*), were expressed at significantly higher levels in R3 KO mice relative to WT (**Figure 1e-g**). These animal experiments indicate that CYB5R3 deficiency in endothelial cells exacerbates LPS induced inflammatory activation, endothelial dysfunction, and systemic hypotension.

**Figure 1.**
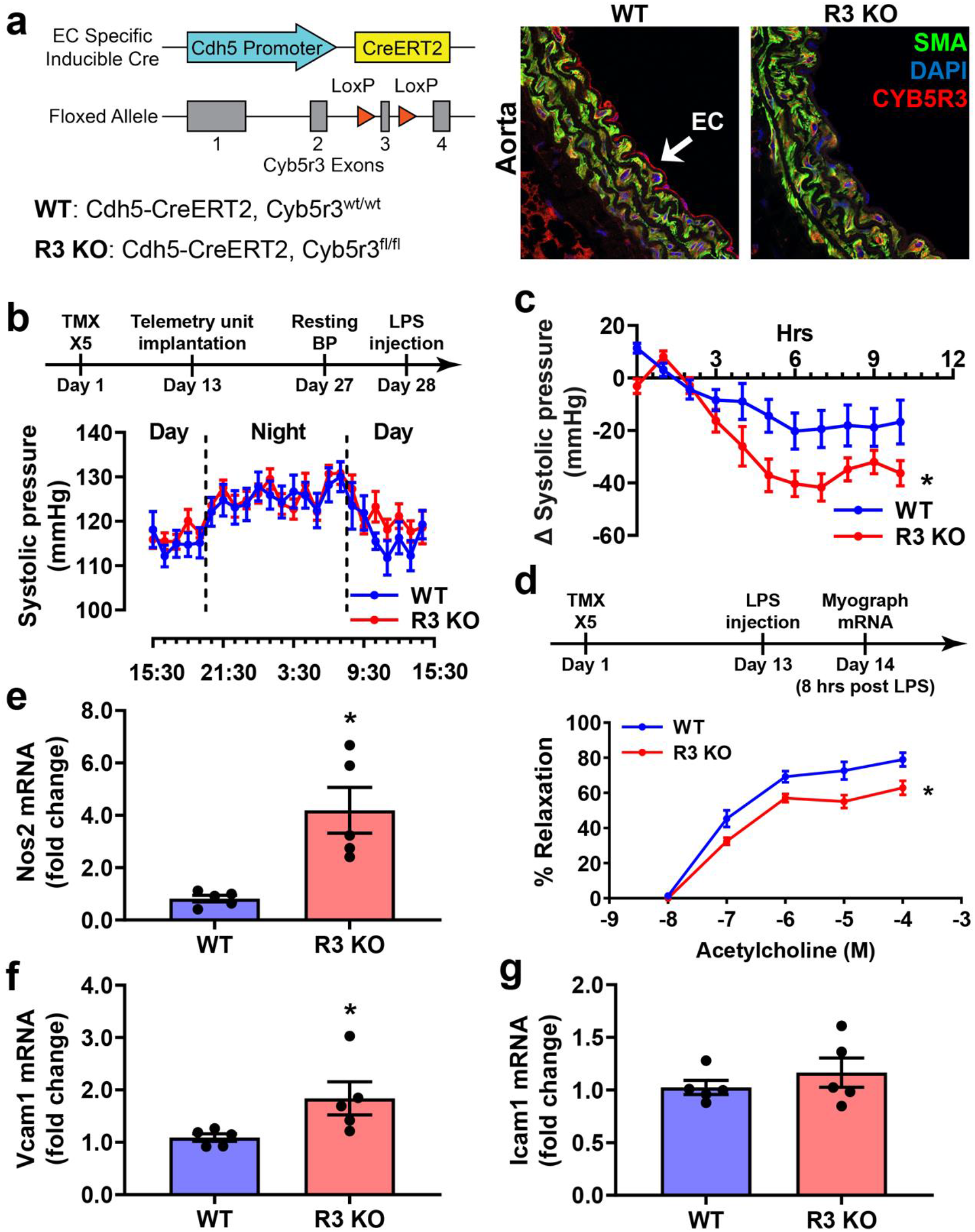
LPS induced vascular dysfunction is exacerbated in endothelium-specific *Cyb5r3* knockout mice. **(a)** *Cdh5-CreER^T^*^2^*/Cyb5r3^wt/wt^* (WT) and *Cdh5- CreER^T^*^2^*/Cyb5r3^fl/fl^* (R3 KO) mice were treated with tamoxifen. Endothelial CYB5R3 expression was examined by immunofluorescence staining in the aorta. **(b)** Mice received five daily tamoxifen (10 mg/kg) injections and subsequent radiotelemetry unit implantation as indicated in the timeline. Systolic pressure was monitored using radiotelemetry for 24 hours immediately before LPS treatment. N= 8 (WT) and 14 (R3 KO). **(c)** LPS (5 mg/kg) intraperitoneal injections stimulated hypotension in both WT and R3 KO mice. By comparing pre- and post-LPS pressures at similar times of the day, the change in systolic pressure was significantly greater in R3 KO mice relative to WT. N= 8 (WT) and 14 (R3 KO); * indicates p <0.05 between WT and R3 KO with 2-way ANOVA. **(d)** In a different cohort, tamoxifen (10 mg/kg) treated mice were injected intraperitoneally with LPS (5 mg/kg) as indicated in the timeline. Aortae were isolated from a different cohort of mice 8 hours after LPS (5 mg/kg) intraperitoneal injections. In *ex vivo* wire myography, R3 KO aortae showed less vasorelaxation with cumulative doses of acetylcholine when compared to WT. N=5; * indicates p<0.05 with 2-way ANOVA. **(e-g).** The mRNA expression levels of *Nos2*, *Vcam-1*, and *Icam-1* were examined in the same aorta samples. N=5; * indicates p<0.05 between WT and R3 KO with unpaired parametric t-test.

To examine whether endothelial CYB5R3 deficiency is directly responsible for the enhanced LPS signaling, we challenged cultured human aortic endothelial cells (HAEC) with LPS overnight. Using 1 μg/ml of LPS, we observed significantly higher VCAM-1 and ICAM-1 protein expression in *Cyb5r3* siRNA-treated HAECs (**Figure 2a-c**). It is known that nuclear factor kappa B (NF-κB) is a major transcription factor responsible for VCAM-1 and ICAM-1 expression^37^. A time course of LPS treatment showed over-activation of NF-κB at the one-hour time point, indicated by augmented phosphorylation of the p65 subunit and enhanced VCAM-1 and ICAM-1 expression at the 4-hour time (**Figure 2d-g**). Moreover, BMS 345541, an inhibitor targeting inhibitor of nuclear factor- κB kinase (IKK) prevented VCAM-1 and ICAM-1 upregulation in *Cyb5r3* knockdown cells, indicating the overactivated inflammatory signaling was NF-κB dependent. Together, these data indicate that CYB5R3 is critical for LPS induced endothelial inflammatory activation both *in vivo and in vitro*.

**Figure 2.**
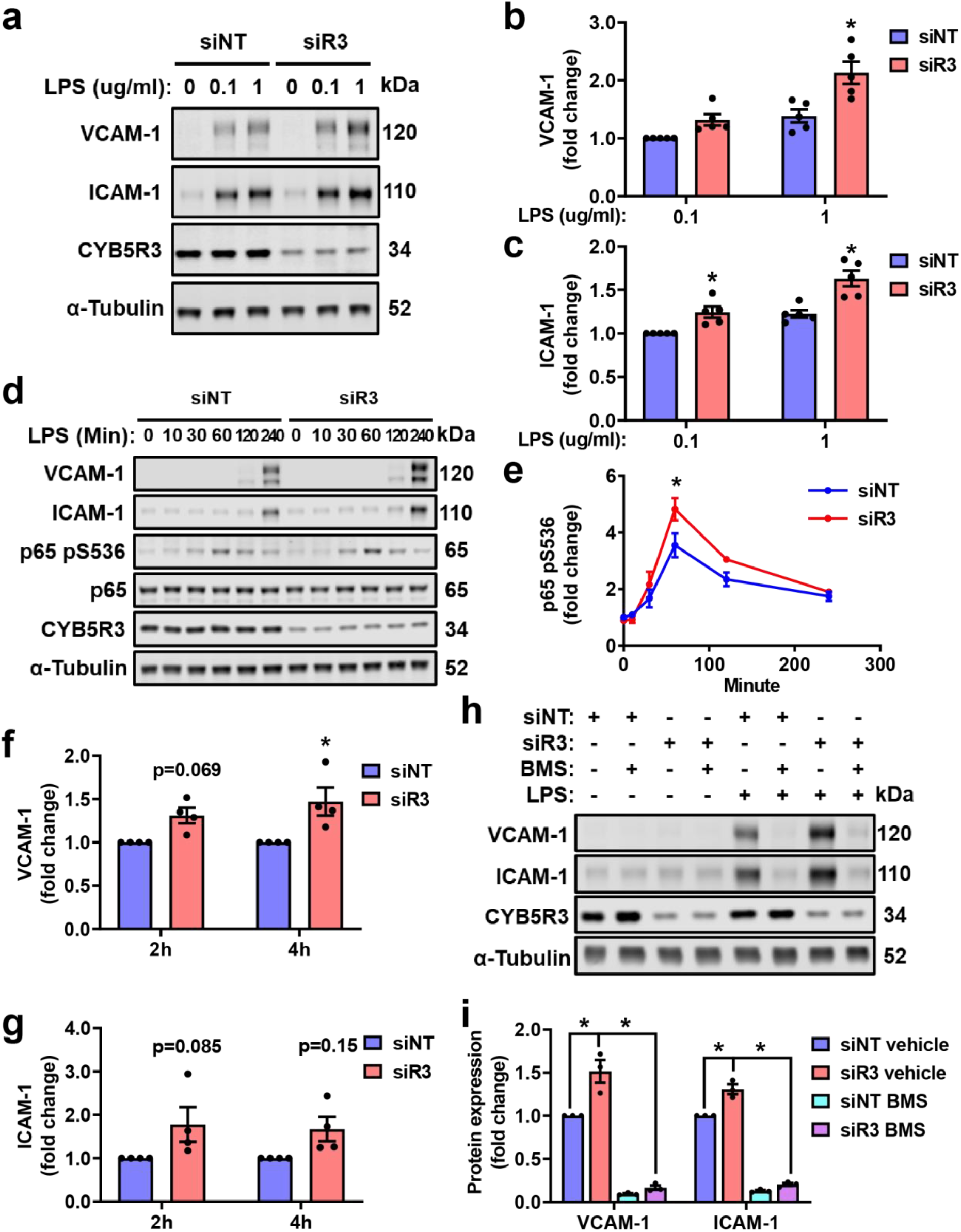
*Cyb5r3* gene silencing potentiates LPS induced NF-kB activation and VCAM-1 and ICAM-1 expression in endothelial culture. HAECs were transfected with non-targeting (siNT) and *Cyb5r3*-targeting (siR3) siRNA for 48 hours before challenge with LPS. **(a-c)** HAECs were treated with 0.1 or 1 µg/ml LPS for 16 hours. At the 1 µg/ml concentration, both LPS-induced VCAM-1 and ICAM-1 protein expressions were significantly higher in the siR3 group. N=5; * indicates p<0.05 between siNT and siR3 with 2-way ANOVA and Sidak post-hoc test. **(d-g)** Cells were treated with 1 µg/ml LPS for up to four hours. VCAM-1 was increased to a significantly higher level in siR3 treated cells relative to siNT and was accompanied by more p65 phosphorylation on S546. N=4; * indicates p<0.05 between siR3 and siNT with 2-way ANOVA and Sidak post-hoc test. **(h-i)** Cells were pretreated with BMS 345541 (10 µM) for 20 minutes and challenged with LPS (1 µg/ml) overnight. N=3; * indicates p<0.05 between linked groups with 2-way ANOVA and Sidak post-hoc test.

### CYB5R3 mitigates LPS signaling in a NOX4 dependent manner

It is well established that LPS signals through toll-like receptor-4 (TLR-4) to elevate ICAM and VCAM-1 expression^38^. Additionally, NOX4 has been shown to mediate LPS- TLR4 signaling in endothelial cells^39^. Considering the regulatory effect of NQO1, a CoQ reductase, on NOX4, we tested whether CYB5R3 modulates NOX4 activity to affect LPS-induced inflammatory activation using cultured HAECs treated with siRNA targeting *Cyb5r3* and *Nox4* mRNA, separately or in combination. Non-targeting siRNA was added as a control for the total siRNA load. As shown in **Figure 3a, 3d, and 3e**, both CYB5R3 and NOX4 protein expression were significantly suppressed by the corresponding siRNA. Interestingly, *Nox4* siRNA slightly but significantly increased CYB5R3 protein expression (**Figure 3d**), suggesting a potential compensatory effect between NOX4 and CYB5R3. More importantly, the loss of NOX4 completely prevented the upregulation of LPS-induced VCAM-1 and ICAM-1 expression in *Cyb5r3* deficient HAECs (**Figure 3a-c**). To examine whether the protective effects of *Cyb5r3* and *Nox4* occur at the transcriptional level, we measured mRNA expression at 2 and 4 hours after LPS treatment. Indeed, both LPS-stimulated *Vcam-1* and *Icam-1* mRNA were increased in *Cyb5r3* knockdown HAECs at the 4-hour time point (**Figure 4a-b**). When HAECs were treated with *Nox4* siRNA, augmentation of *Vcam-1* mRNA in the *Cyb5r3* deficient HAECs was ablated (**Figure 4a**). By contrast, *Nox4* gene silencing did not significantly affect *Icam1* mRNA at either time point (**Figure 4b**). It is possible that LPS-induced *Vcam-1* and *Icam-1* expression follow different time courses. Knockdown efficiencies of *Cyb5r3* and *Nox4* were verified at the mRNA level (**Figure 4c-d**). Surprisingly, the compensatory upregulation of CYB5R3 protein in *Nox4* knockdown cells was not significantly reflected at the mRNA level (**Figure 4c**). However, *Nox4* mRNA was significantly increased with *Cyb5r3* gene silencing (**Figure 4d**).

**Figure 3.**
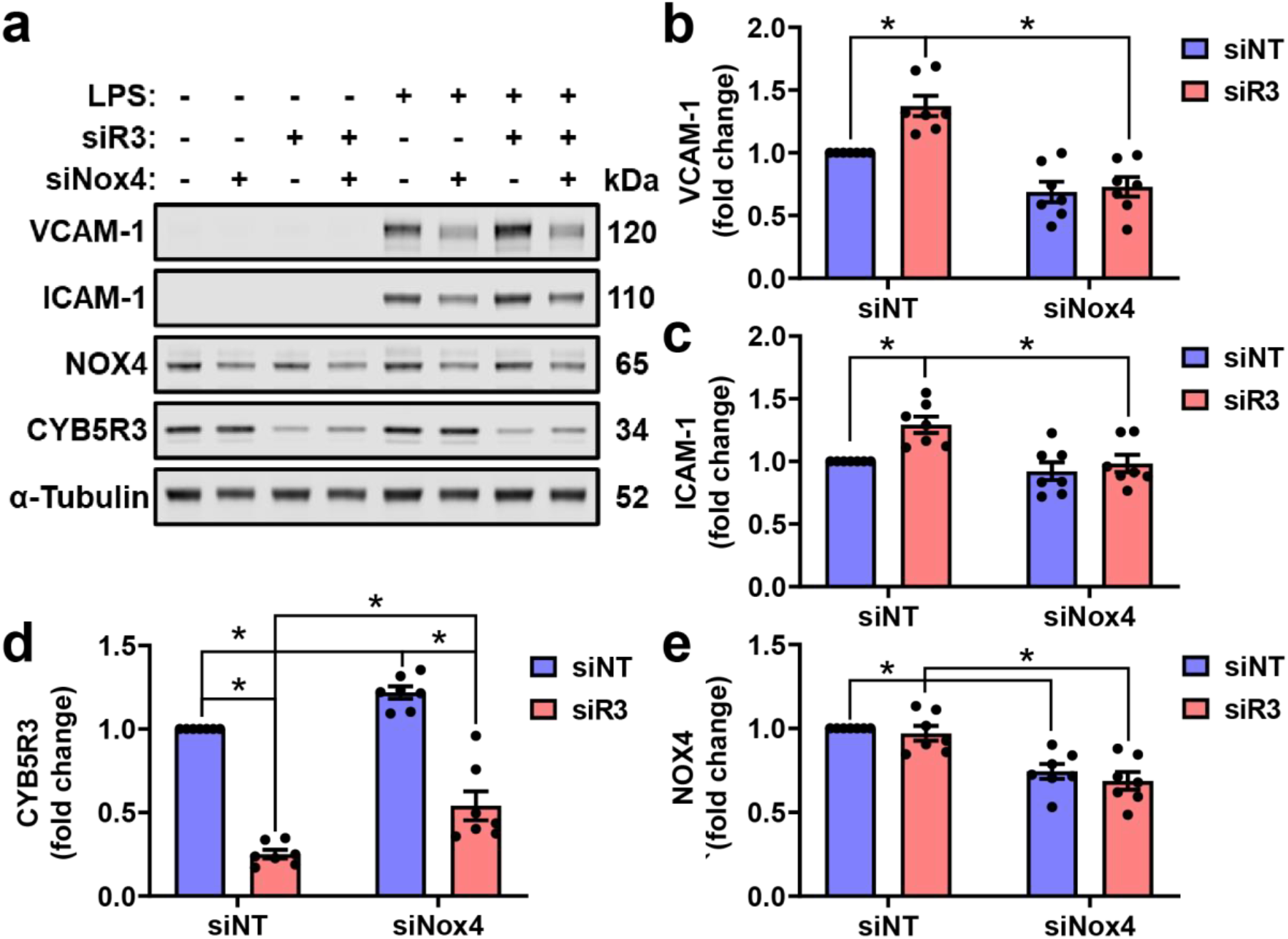
The anti-inflammatory effects of CYB5R3 are NOX4 dependent. **(a)** HAECs were co-transfected with siRNA targeting *Cyb5r3* (siR3) and *Nox4* (siNox4); non-targeting siRNA (siNT) was used to keep the siRNA load the same between groups. At 48 hours following transfection, cells were treated with 1 µg/ml LPS for 16 hours. **(b-e)** Protein expression levels of VCAM-1, ICAM-1, CYB5R3, and NOX4 were quantified at the 16-hour time point. N=7; * indicates p<0.05 between linked groups with 2-way ANOVA and Tukey’s multiple comparisons.

**Figure 4.**
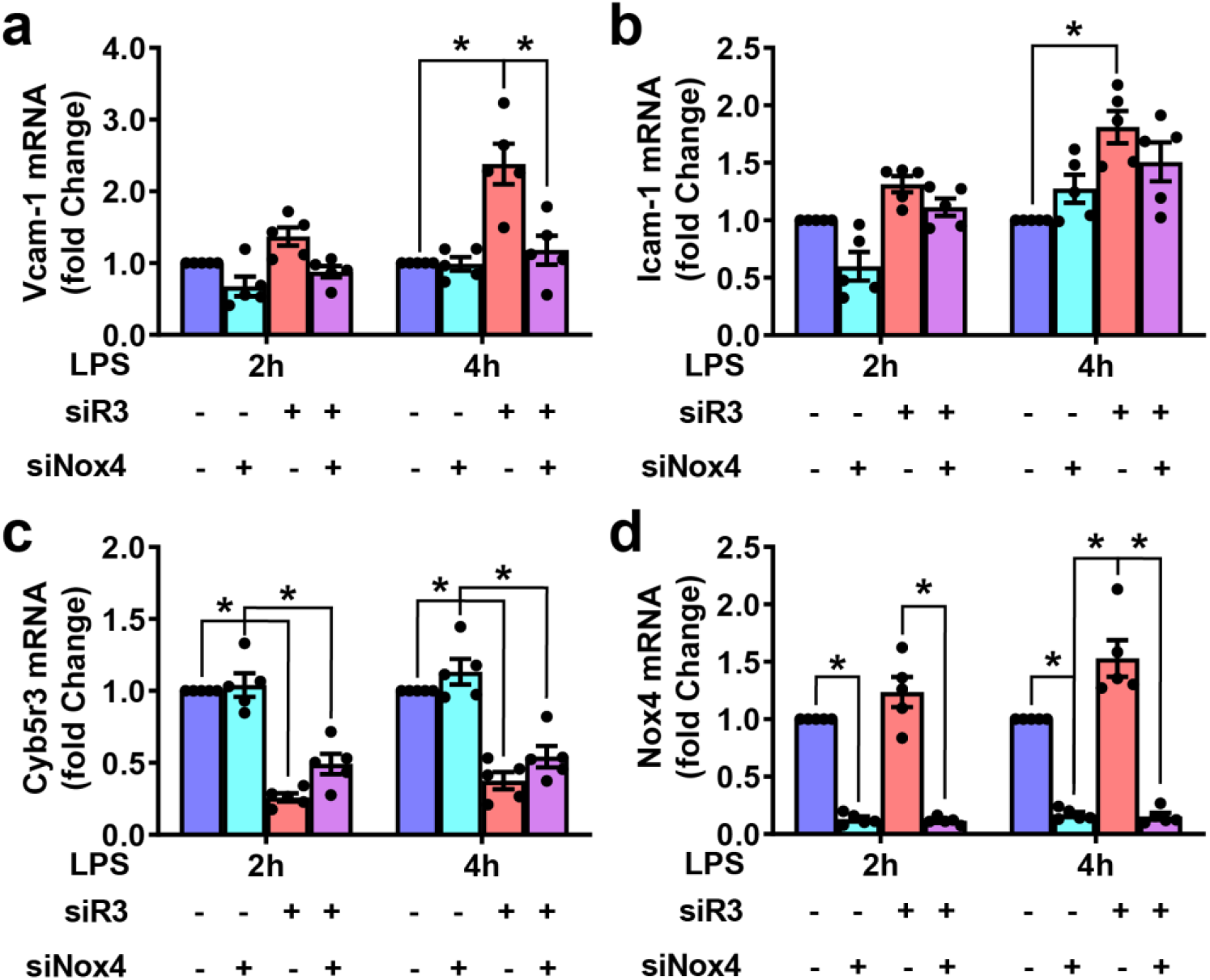
NOX4 regulates *Vcam1* and *Icam1* expression at the transcript level. HAECs were co-transfected with siRNA targeting *Cyb5r3* (siR3) and *Nox4* (siNox4); non-targeting siRNA (siNT) was used to keep the siRNA load the same between groups. 48 hours after the siRNA transfection, cells were treated with 1 µg/ml LPS and collected 2 or 4 hours after the LPS treatment to examine mRNA expression. N=5; * indicates p<0.05 between linked groups with 2-way ANOVA and Tukey’s multiple comparisons.

### CYB5R3 and NOX4 modulate TNF-α induced VCAM-1 expression

LPS-induced inflammatory signaling *in vivo* can be magnified and propagated via TNF- α^40, 41^. Therefore, we extended our studies to test if the protective effects of CYB5R3 involved TNF-α signaling. In cultured HAECs, in which *Cyb5r3* was silenced using siRNA prior to overnight treatment with TNF-α (10 ng/ml), Cyb5R3 deficiency exacerbated both VCAM-1 and ICAM-1 protein expression. When *Nox4* was silenced alongside *Cyb5r3*, only the previously seen VCAM-1 upregulation with *Cyb5r3* deficiency was prevented, not ICAM-1. (**Figure 5a-c**). Therefore, CYB5R3 protects, in part, against TNF-α mediated inflammatory signaling. Moreover, we observed the same compensatory effects on CYB5R3 and NOX4 expression (**Figure 4d-e, 5d-e**).

**Figure 5.**
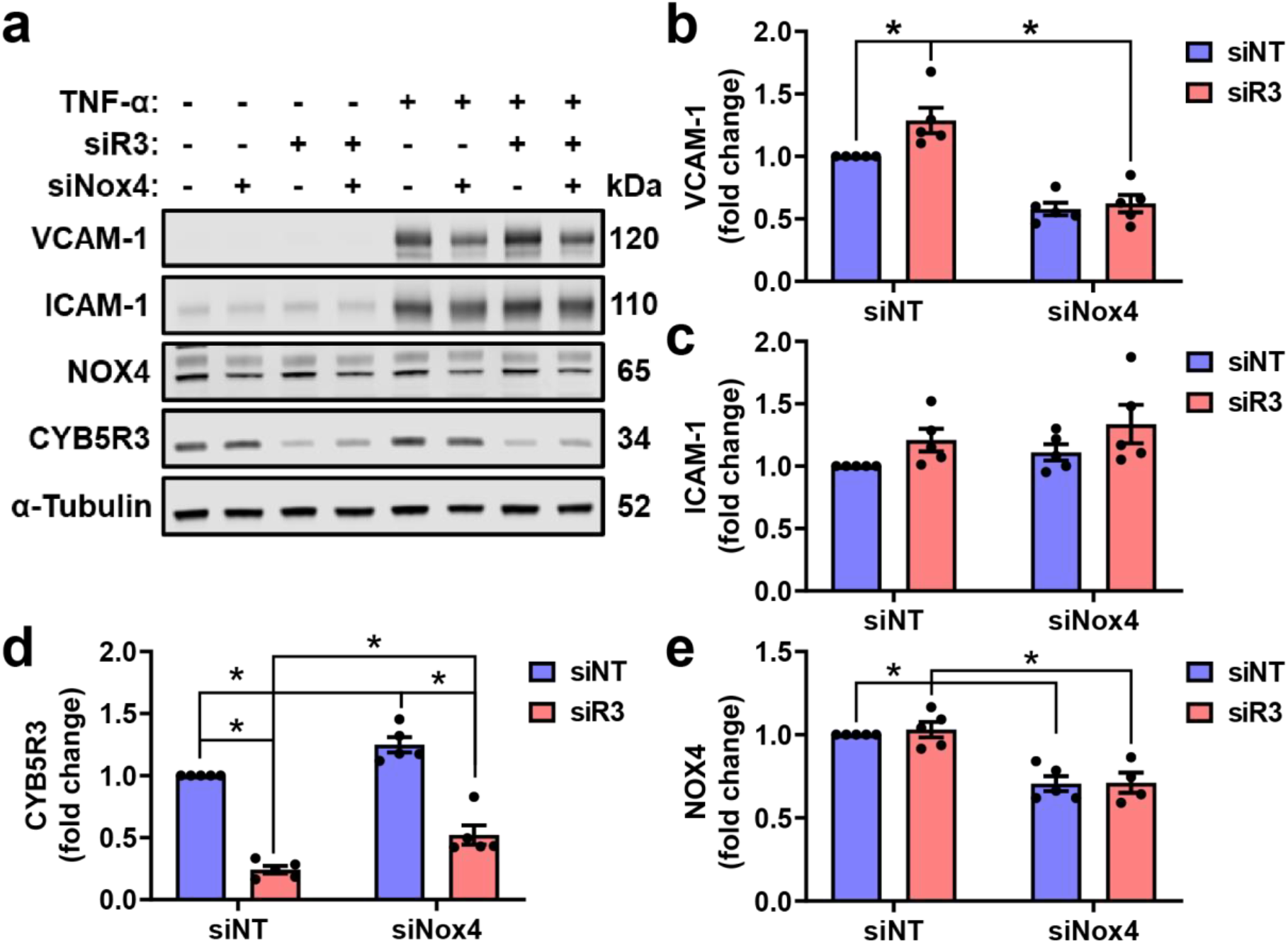
Endothelial CYB5R3 and NOX4 regulate TNF-α induced inflammatory signaling in a similar pattern. **(a)** HAECs were co-transfected with non-targeting (siNT), *Cyb5r3-*targeting (siR3), and *Nox4*-targeting (siNox4) siRNA as indicated. TNF-α was added to the medium 48 hours after transfection, and cells were collected after 16 hours for immunoblotting. **(b-e)** Protein expression levels of VCAM-1, ICAM-1, CYB5R3, and NOX4 were quantified at the 16-hour time point. N=7; * indicates p<0.05 between linked groups with 2-way ANOVA and Tukey’s multiple comparisons.

### CYB5R3 and NOX4 interact on the mitochondrial outer membrane

Since our data indicated a protective role for CYB5R3 in LPS and TNF-α-activated endothelial inflammation mediated by NOX4, we hypothesized that CYB5R3 may regulate NOX4 activity through ubiquinone reduction on the mitochondrial membrane. Mitochondria are a primary location for both CYB5R3 and NOX4^11, 42–44^. To test this hypothesis, we first performed immunostaining for CYB5R3 and NOX4 in cultured HAECs. Both CYB5R3 and NOX4 colocalized with TOM20, a mitochondrial marker (**Figure 6a**). Their mitochondrial localization was better demonstrated by the super-resolution images stained with the same antibodies (**Figure 6b**).

**Figure 6.**
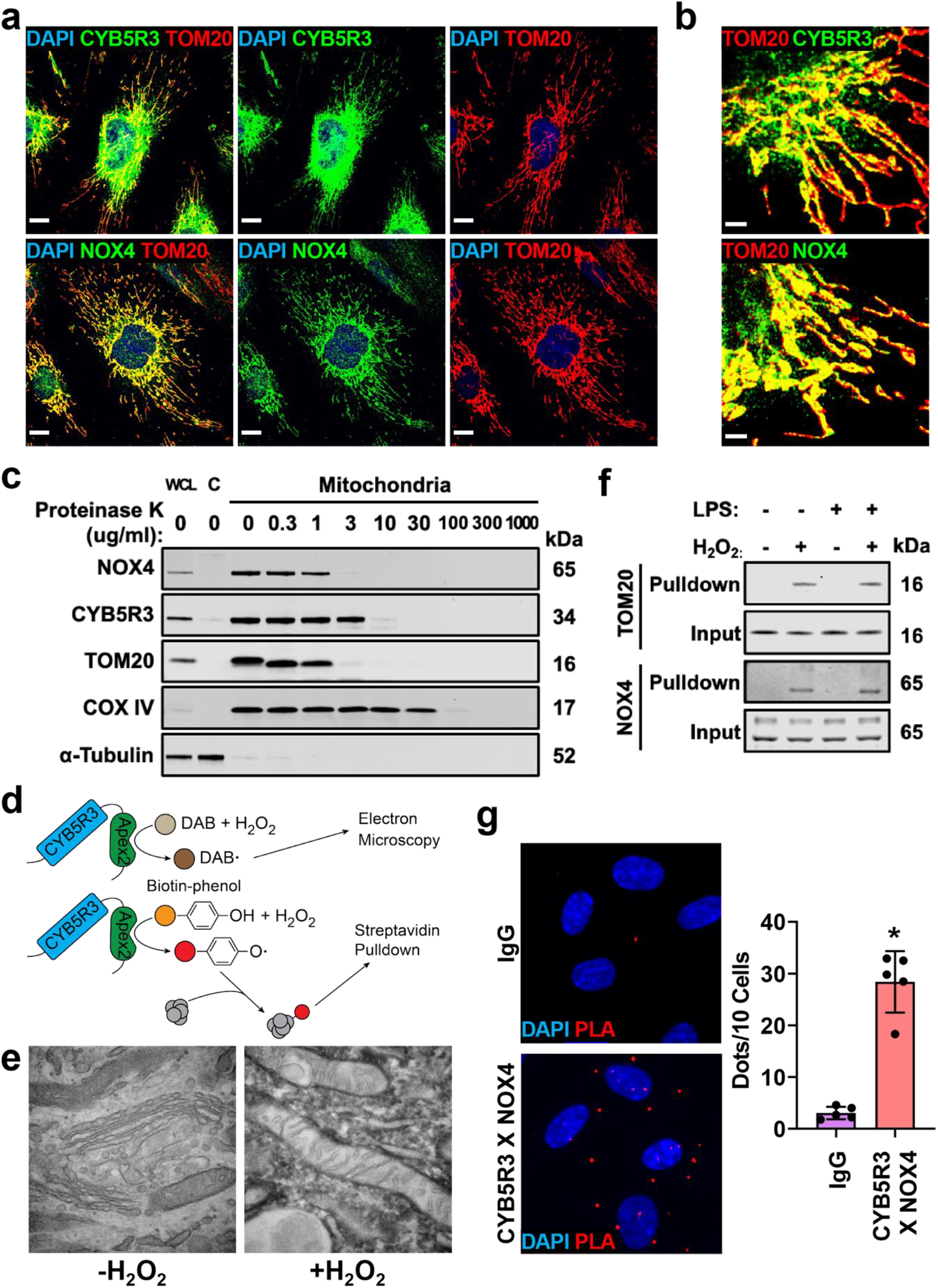
CYB5R3 resides in close proximity to NOX4 in mitochondria. **(a)** Confluent HAECs were stained with CYB5R3 or NOX4 (green), TOM20 (red), and DAPI (blue). Both CYB5R3 and NOX4 colocalize with TOM20 in confocal images. The scale bar represents 10 µm. **(b)** The mitochondrial localization of CYB5R3 and NOX4 is shown in super-resolution images (STED). The scale bar represents 2 µm. **(c)** Mitochondria were isolated from HEK293FT cells and digested in various concentrations of proteinase K. Proteins on the outer membrane were more susceptible to digestion compared to inner membrane proteins. **(d)** Strategy using CYB5R3-APEX2 to study protein subcellular localization and protein-protein interactions. **(e)** The electron microscopic image shows the electron-dense area around mitochondria in CYB5R3- APEX2 expressing HAECs only in the presence of H_2_O_2_. **(f)** Biotinylated proteins were captured from CYB5R3-APEX2 expressing HAECs. In the presence of H_2_O_2_, both TOM20 and NOX4 were detected in the pulldown samples irrespective of LPS treatment (1 µg/ml for one hour). **(g)** The proximity ligation assay (PLA, red) confirmed direct interactions between CYB5R3 and NOX4 (CYB5R3 X NOX4). The PLA positive signals were quantified as dots per 10 nuclei (DAPI, blue). The quantification represents average count ± standard deviation from 5 images in each group. * indicates p<0.05 with Student’s t-test.

Moreover, we sought to determine whether CYB5R3 and NOX4 were expressed on the outer or inner mitochondrial membrane by exposing isolated mitochondria to proteinase As shown in **Figure 6c**, CYB5R3 and NOX4 were fully digested at 10 and 3 ug/ml proteinase K, respectively, which similarly occurred with the mitochondrial outer membrane protein TOM20 (**Figure 6c**). In comparison, cytochrome c oxidase subunit 4 (COX IV), an inner membrane protein, was resistant to proteinase K (30 μg/ml). These findings indicated that CYB5R3 and NOX4 reside on the mitochondrial outer membrane.

Additional studies were performed to investigate CYB5R3’s spatial localization and protein-protein interactions. APEX2, a modified soybean peroxidase used to examine subcellular localization and protein interacting partners, was fused to the C-terminus of CYB5R3 (**Figure 6d**)^29, 30^. In the presence of hydrogen peroxide (H_2_O_2_), CYB5R3- APEX2 converts 3,3′-Diaminobenzidine into brown precipitates that can be visualized in electron microscopy as an electron-dense signal. In HAECs overexpressing CYB5R3- APEX2, the electron-dense signal accumulated in the outer periphery of mitochondria, suggesting CYB5R3 resides on the outside of mitochondria (**Figure 6e**), consistent with the long-standing hypothesis on CYB5R3’s orientation on the mitochondrial outer membrane^45^. Since APEX2 can be activated by high concentrations of H_2_O_2_ to biotinylate proximal proteins with biotin-phenoxyl radical, we also used it to pull down biotinylated proteins from CYB5R3-APEX2 expressing HAECs. Our pulldown results included TOM20 and NOX4 (**Figure 6f**), suggesting both proteins are in close proximity to CYB5R3. Proximity ligation assay was also used and revealed endogenous CYB5R3 and NOX4 to be spatially close to each other in endothelial cells (**Figure 6g**).

### Membrane-bound CYB5R3 regulates NOX4 activity

Since CYB5R3 and NOX4 reside in close proximity to each other on the mitochondrial outer membrane, we hypothesized that CYB5R3 interacts with NOX4 to regulate its activity. NOX4 is known to produce predominantly H_2_O_2_ and a portion of O_2_^•-^^12^. HAECs treated with *Cyb5r3* siRNA showed decreased H_2_O_2_ production and increased mitochondrial O_2_^•-^ (**Figure 7a-b**). Electron leakage from mitochondrial respiration is a major source of O_2_^•-^. Therefore, we examined mitochondrial function to distinguish the source of reactive oxygen species in *Cyb5r3* knockdown HAECs. The results showed that the lack of CYB5R3 did not affect protein expression levels of respiration complexes (**Supplementary Figure 2)**, mitochondrial respiration (**Supplementary Figure 3a-g**), or mitochondrial membrane potential (**Supplementary Figure 3h**) with or without LPS stimulation. Furthermore, by quantifying the superoxide specific DHE oxidation product, 2OHE^+^, using HPLC-MS, we confirmed increased superoxide anion levels in *Cyb5r3* knockdown HAECs was NOX4 dependent (**Figure 7c**). To better understand how CYB5R3 affects NOX4 activity, we created expression vectors carrying wild-type CYB5R3, inactive CYB5R3 (K111A or K111A/G180V), and non-membrane bound active CYB5R3 missing the N-terminal membrane anchor (ΔAA1-23). To exclude the effect of endogenous CYB5R3, we generated *Cyb5r3* deficient HEK293FT cell lines using lentiviral shRNA (**Supplementary Figure 3a**). CYB5R3 activity was verified in *Cyb5r3* deficient HEK293FT cells forced to express similar amounts of mutant CYB5R3 (**Figure 7d**). Additionally, to test NOX4 activity, *Cyb5r3* deficient HEK293FT cells were co-transfected with NOX4 and its cofactor p22 (**Supplementary Figure 3b**). These experiments revealed significantly increased H_2_O_2_ produced from NOX4 in cells with wild-type CYB5R3, while inhibitory effects were observed in cells transfected with inactive CYB5R3. (**Figure 7e**). The activity of NOX4 did not appear to be affected in cells transfected with the non-membrane bound active CYB5R3 (ΔAA 1-23), which lacks membrane binding. (**Figure 7e**). These data suggest that both the activity and membrane localization of CYB5R3 are required for NOX4 to produce H_2_O_2_.

**Figure 7.**
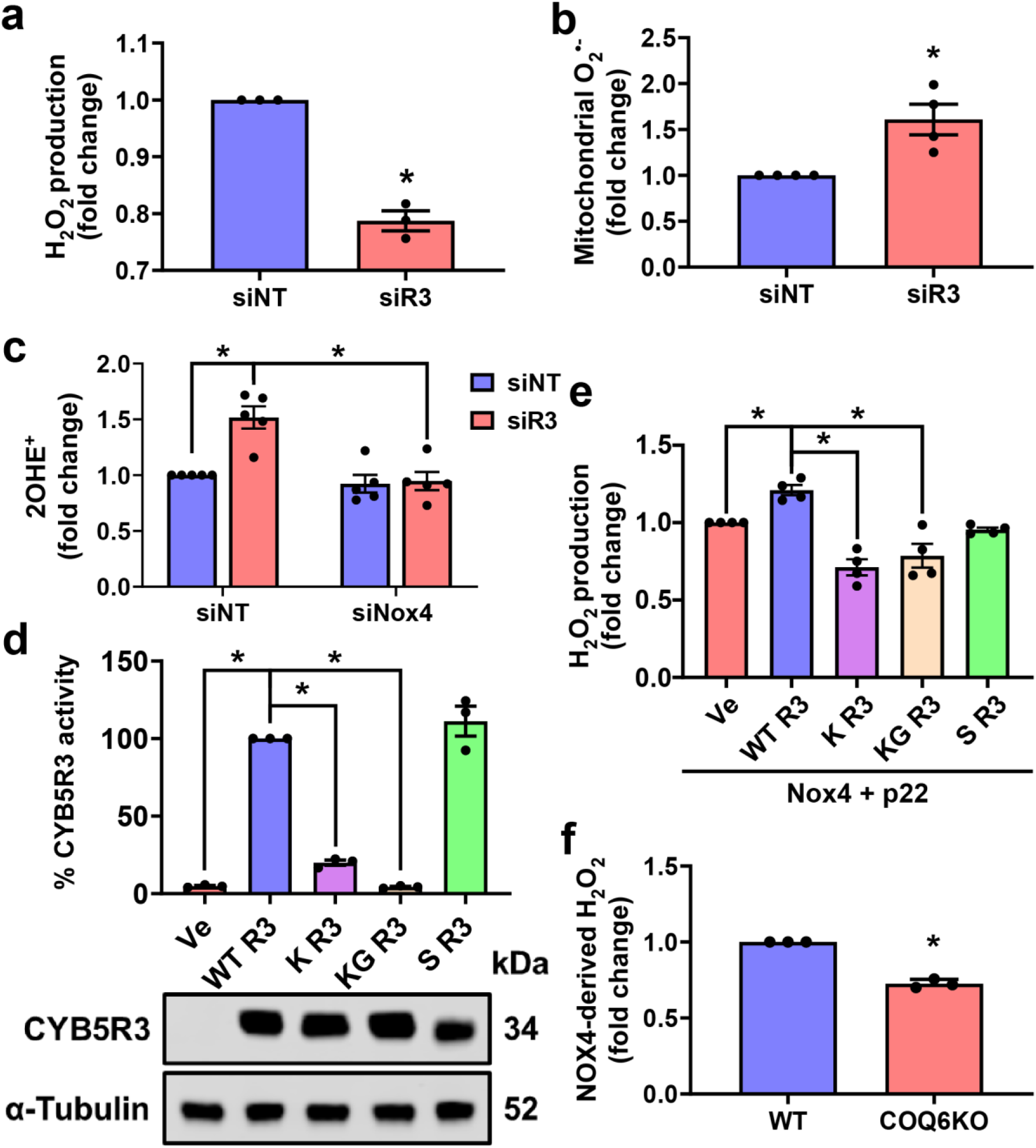
Optimal NOX4 production of H_2_O_2_ requires CYB5R3 membrane localization and activity. (a-c) HAECs were transfected with non-targeting (siNT), *Cyb5r3*-targeting (siR3), and Nox4-targeting (siNox4) siRNA for 48 hours. **(a-b)** H_2_O_2_ and mitochondrial O_2_^•-^ were measured using coumarin boronic acid and MitoSOX, respectively. N=3-4; * indicates p<0.05 between siNT and siR3 with ratio paired t-test. **(c)** 2OHE^+^ was quantified with HPLC-MS. N=5; * indicates p<0.05 between linked groups with 2-way ANOVA and Tukey’s multiple comparisons. **(d)** *Cyb5r3* knockdown HEK293FT cells were forced to express wild-type CYB5R3 (R3 WT), K111A CYB5R3 (R3 K111A), K111A/G180V CYB5R3 (R3 K111A/G180V), and non-membrane bound active CYB5R3 (R3 ΔAA1-23) to similar levels. CYB5R3 activities were measured with the pseudosubstrate 2,6-Dichlorophenolindophenol (DCPIP). N=3; * indicates p<0.05 between indicated groups with one-way ANOVA and Dunnett’s multiple comparisons. **(e)** NOX4 activities were evaluated by the amount of H_2_O_2_ produced from NOX4 and p22 expressing HEK293FT cells. NOX4 derived H_2_O_2_ varied between HEK293FT cells with different forms of coexpressed CYB5R3. N=4; * indicates p<0.05 between indicated groups with one-way ANOVA and Dunnett’s multiple comparisons. **(f)** Wildtype (WT) and *COQ6* knockout (COQ6KO) HEK293 cells were forced to express NOX4 and p22, and H_2_O_2_ production was measured. N=3; * indicates p<0.05 with ratio paired t-test.

### Endogenous CoQ plays a role in NOX4-dependent H_2_O_2_ production

To test if the regulation of CYB5R3 on NOX4 activity is mediated by CoQ, we measured H_2_O_2_ generated from NOX4 in *COQ6* knockout HEK293 cells unable to synthesize endogenous CoQ^28^. Consistent with our hypothesis, NOX4’s activity to generate H_2_O_2_ was compromised in CoQ deficient cells (**Figure 7f**). Since catalase expression was higher in the *COQ6* knockout HEK293 cells than the wild-type control cells (**Supplementary Figure 4a-c)**, we tested whether the apparent loss of H_2_O_2_ production from NOX4 was due to greater H_2_O_2_ consumption by catalase. The use of 1,3- aminotriazole (ATZ), a catalase inhibitor, failed to increase H_2_O_2_ steady-state levels arising from NOX4 in *COQ6* knockout cells, indicating that the compromised phenotype was most likely independent of catalase activity (**Supplemental Figure 4d**). Our data confirmed that endogenous CoQ is required for optimal NOX4-dependent H_2_O_2_ production.

## DISCUSSION

Vascular wall inflammation is allied to an imbalance of reduction-oxidation (redox) signaling. However, the fundamental mechanisms that modulate redox homeostasis in the setting of vascular inflammation are not fully understood. In this study, we uncover significant breakthroughs that advance the understanding of redox signaling and endothelial inflammatory activation. First, by generating a novel endothelium-specific CYB5R3 knockout mouse, we discover that knockout animals subjected to endotoxin exhibit exacerbated hypotension, endothelial cell dysfunction, and increased vascular wall inflammation, identifying a previously unrecognized role of CYB5R3 in the endothelium. Second, we discover for the first time that CYB5R3 regulates inflammation through an NADPH oxidase 4 (NOX4) -dependent mechanism to facilitate the formation of H_2_O_2_. Lastly, we show that mitochondrial membrane-bound CYB5R3 regulates NOX4’s ability to produce hydrogen peroxide via endogenous CoQ. Collectively, these new findings highlight the functional significance of endothelial CYB5R3 in vascular wall inflammation and a previously unrecognized partnership between CYB5R3 and NOX4 via CoQ to produce H_2_O_2_ in order to control inflammatory signaling (**Figure 8**).

**Figure 8.**
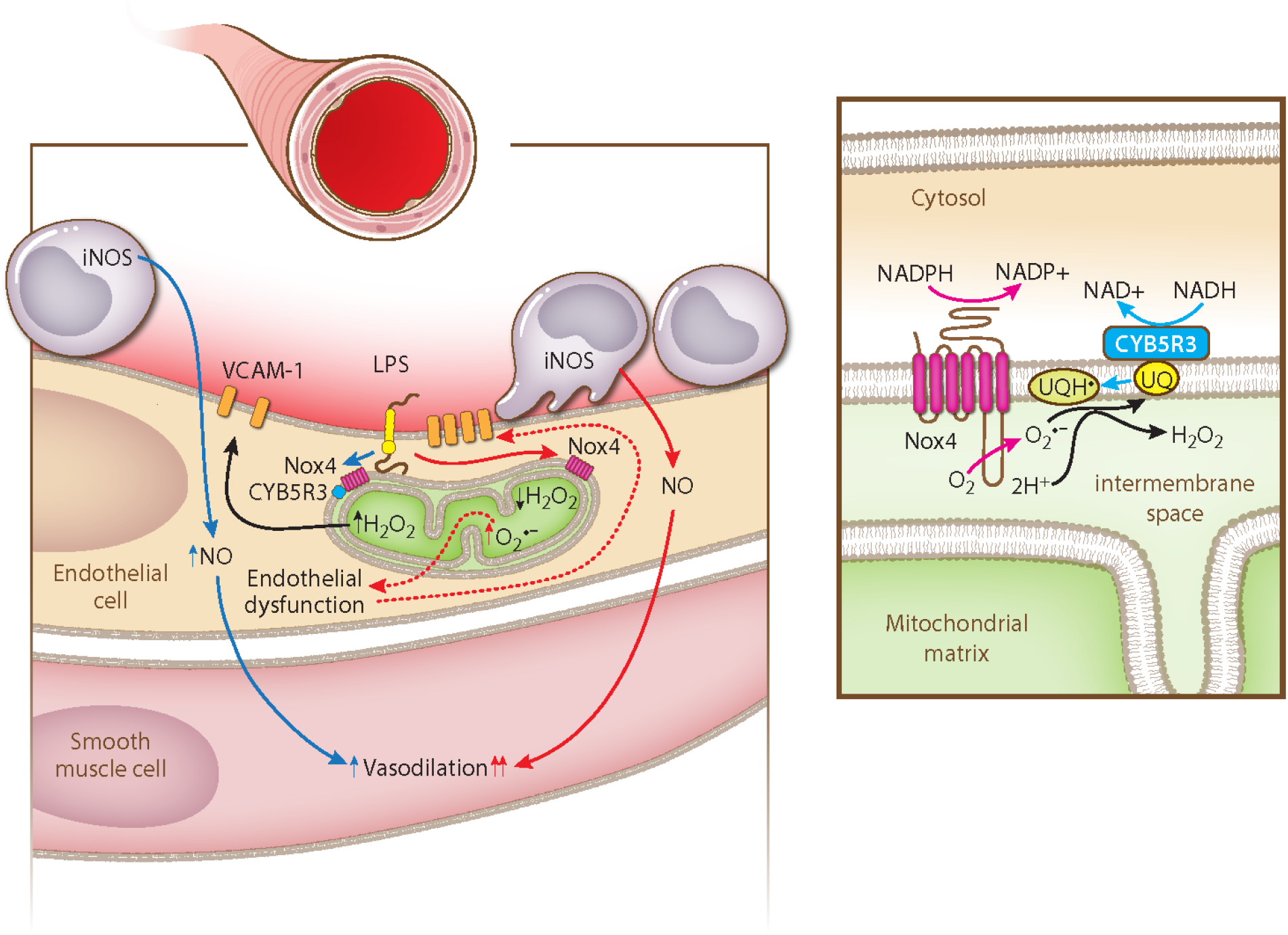
Schematic for the regulation of CYB5R3 on endothelial inflammatory activation and NOX4 activity. In the right panel, CYB5R3 reduces the fully oxidized coenzyme Q, ubiquinone (UQ), to semiubiquinone (UQH^•^) at the expense of NADH. Meanwhile, NOX4 reduces a molecule of oxygen (O_2_) to superoxide (O_2_^•-^) in the mitochondrial intermembrane space. The resultant O_2_^•-^ and UQH^•^ react to generate H_2_O_2_ and UQ. CYB5R3 normally facilitates NOX4 predominant production of H_2_O_2_ in mitochondria (left panel). During endothelial inflammatory activation, *e.g.,* with LPS challenge, H_2_O_2_ derived from NOX4 upregulates VCAM-1 expression on the endothelial surface to recruit monocytes. NOS2 in recruited monocytes produces nitric oxide (NO), causing extensive vasodilation and systemic hypotension. When CYB5R3 activity is impaired in endothelial cells (right panel), NOX4 produces less H_2_O_2_ and more O_2_^•-^, which results in elevated VCAM-1 expression, enhanced monocyte recruitment, more NOS2 derived NO, and more severe hypotension.

NOX4 is a member of the family of NADPH oxidases that reduces O_2_ to reactive oxygen species. Among its family members, NOX4 is unique as it is constitutively active and generates predominantly H_2_O_2_. Different mechanisms have been proposed to explain NOX4’s ability to produce H_2_O_2_ rather than O_2_^•-^^10, 12^. One theory is that NOX4 facilitates the dismutation of newly formed O_2_^•-^ to H_2_O_2_ (Eq 1)^10^. On the plasma membrane, NOX4 is a six-transmembrane protein with both *N* and *C*-termini in the cytosol^46^. Among NOX4’s three extracellular loops, the third loop is 28 amino acids longer than the corresponding loops in NOX1 and NOX2 that are dedicated to O_2_^•-^ production. Takac and colleagues demonstrated that NOX4, lacking the third extracellular loop or with substitution of two essential cysteines in that loop (Cys 226 and Cys270), switched to preferentially producing O_2_^•-^ versus H_2_O_2_^10^. The authors hypothesized that Cys226 and Cys270 may form a disulfide bond to retain a newly formed O_2_^•-^, increasing the rate of spontaneous dismutation into H_2_O_2_. Moreover, the authors suggested that His222 in this loop helps donate protons to the retained O_2_^•-^.

Contradictory to the hypothesized intrinsic dismutase activity of NOX4, breaking the Cys226-Cys270 by β-mercaptoethanol enhanced NOX4’s ability to produce H_2_O_2_^18^. Interestingly, NOX4 possesses a putative quinone binding site (AA203-209) that can be used to interact with CoQ. Exposing NOX4 to certain quinone compounds or NQO1, a major CoQ reductase, induced H_2_O_2_ production from NOX4. Furthermore, Nisimoto Y. *et al.* analyzed the kinetics of the H_2_O_2_ formation by NOX4, which did not support the intrinsic dismutase activity. Instead, their data suggested a one-electron reduction of O_2_^•-^ to H_2_O_2_ (Eq 2)^12^.

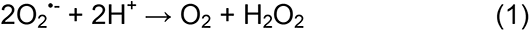

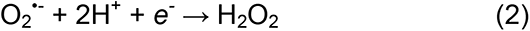

In our results, endothelial cells with *Cyb5r3* gene silencing had decreased H_2_O_2_ and increased O_2_^•-^ production (**Figure 7a-b**). Since mitochondrial respiration was not affected by knocking down *Cyb5r3*, the elevated O_2_^•-^ was not likely due to electron leakage from the respiratory chain (**Supplementary Figure 1-2**). Indeed, simultaneously silencing *Nox4* prevented O_2_^•-^ from *Cyb5r3* knockdown endothelial cells (**Figure 7c**). Furthermore, NOX4-derived H_2_O_2_ can be promoted by coexpressing the wild-type CYB5R3 but is suppressed when inactive mutants are expressed (**Figure 7e**). Additionally, NOX4 activity was not affected by the enzymatically active CYB5R3 missing its membrane binding capacity (**Figure 7e**). These findings pointed to a crucial role for both the activity and subcellular location of CYB5R3 in the optimal function of NOX4. Therefore, we hypothesized that NOX4 generates H_2_O_2_ with O_2_^•-^ reductase activity that was fulfilled at least partially in combination with CYB5R3.

In support of this hypothesis, we found that CoQ deficiency in *COQ6* knockout HEK293 cells resulted in diminished H_2_O_2_ production from NOX4 (**Figure 7f**). To our knowledge, this is the first time that endogenous CoQ has been shown to regulate NOX4 activity, which is in agreement with previous findings for synthetic quinone compounds and NQO1 overexpression^18^. Importantly, although NQO1 is known to regulate mitochondrial function, its localization on the mitochondrial membrane remains controversial^47–51^. While NQO1 can be detected in mitochondria isolated from multiple mouse tissues, it was not found in mitochondria in a human cell line^50^. This disagreement in NQO1’s subcellular localization may be rooted in the difference in species, tissues, and antibody sensitivity. Nonetheless, we clearly demonstrate that CYB5R3 is enriched on the mitochondrial membrane using immunofluorescent imaging and electron microscopy in primary endothelial cells (**Figure 6a-band 6e**), which is consistent with previous reports^42, 43^. Specifically, our data indicate both NOX4 and CYB5R3 colocalize on the mitochondrial outer membrane (**Figure 6c and 6e**). We further show with proximity ligation assay that NOX4 and CYB5R3 are close neighbors, permitting the two enzymes to interact (**Figure 6g**). Therefore, it is possible that CYB5R3, like NQO1, reduces ubiquinone to regulate NOX4 activity.

To further understand the interaction of CYB5R3 and NOX4, the orientation of both enzymes on the mitochondrial outer membrane needs to be defined. Similar to NOX4, CYB5R3 can also be found on the plasma membrane. While the former is transmembrane, the latter a cytosolic protein anchored to the membrane by a short N- terminal sequence^52^. Although we found that endothelial CYB5R3 and NOX4 predominantly reside on the outer mitochondrial membrane, their orientations on the plasma membrane may help with understanding their protein assembly on the mitochondrial membrane. By creating a chimeric CYB5R3 with APEX2 fused to the reductase’s C-terminus, we found that CYB5R3, NOX4, and TOM20 are all in close proximity (**Figure 6f**). APEX2-catalyzed biotin-phenoxyl radicals rapidly react with electron-rich amino acids on proteins in close proximity, 20-nm, because the radicals are short-lived and thus have a short radius of diffusion^53^. Since TOM20 is a transmembrane protein on the mitochondrial outer membrane, our APEX2 data indicate that CYB5R3 and NOX4 are on the outer membrane. Moreover, the biotin-phenoxyl radicals are not membrane permeable, limiting biotinylation of proteins to one side of the membrane, as shown by our electron microscopy image in which APEX2-fused CYB5R3 accumulation occurs outside mitochondria and not in cristae. Therefore, the evidence points to CYB5R3 being on the cytosolic side of the outer membrane. If we can assume that the orientation of CYB5R3 and NOX4 on the mitochondrial outer membrane is the same as on the plasma membrane, NOX4 should have both its termini in the cytosol and its “extracellular loops” in the intermembrane space (**Figure 8**).

The orientation of CYB5R3 and NOX4 is crucial for their collaboration with respect to O_2_^•-^ reduction. As a one-electron reductase, CYB5R3 reduces ubiquinone to semiubiquinone at the expense of NADH. On the mitochondrial outer membrane, CYB5R3 is close to NOX4, allowing physical interaction between them or by association with a third player, ultimately enabling NOX4 to access semiubiquinone produced by CYB5R3. Meanwhile, NOX4 uses NADPH to produce O_2_^•-^ that is transiently trapped at the third “extracellular loop”. With the electron donated by semiubiquinone and the proton donated by His222, O_2_^•-^ is reduced to H_2_O_2_. Furthermore, the high proton concentration in the intermembrane space, maintained by the electron transport chain, can also favor such a reaction.

O_2_^•-^ dismutase (SOD) and reductase (SOR) represent two distinct systems for regulating O_2_^•-^ in cells. Although SORs have been found in lower organisms, their existence in mammalian cells is still under debate^54^. By contrast, three human isoforms of SODs (SOD1-3) have been identified in mammalian cells, having abundant distribution in the cytosol, mitochondria, and extracellular space. Since SOD is extremely effective at converting O_2_^•-^ to H_2_O_2_ (∼10^9^ M^-1^s^-1^)^55, 56^, an immediate question is why human endothelial cells would require an additional SOR to detoxify O_2_^•-^. A closer examination of the SOD isoform subcellular location predicates the necessity for such an alternative reductase system. On the one hand, SOD2 has a mitochondrial targeting sequence that leads it to the mitochondrial matrix^57^. On the other hand, SOD1 is synthesized, matured, and accumulated in the cytosol. Although there is some evidence that in yeast, SOD1 is imported to mitochondria, there is no evidence of SOD1 in mammalian cells translocating to mitochondria unless associated with neurodegenerative disorders^58^. Furthermore, we have confirmed the absence of SOD1 in isolated mitochondria (**Supplementary Figure 4a**). With this current understanding of the SOD isoforms’ subcellular locations, the mitochondrial intermembrane space seems to be devoid of any SOD. Since NOX4 resides on the mitochondrial outer membrane with its critical loop in the intermembrane space, O_2_^•-^ can be trapped there and reduced to H_2_O_2_. It is possible that the semiubiquinone radical produced by CYB5R3, which forms on the cytosolic side of the membrane, flips across the outer mitochondrial membrane to get oxidized by intermembrane O_2_^•-^ to form ubiquinone and H_2_O_2_. This “flip-flop” action of CoQ is fast enough to support a biological function such as the electron transport chain^59^.

The interaction between CYB5R3 and NOX4 has significant implications for endothelial inflammatory activation. *In vitro*, LPS induced expression of VCAM-1 and ICAM-1 was enhanced by the loss of CYB5R3 in endothelial cells on both protein and mRNA levels (**Figure 3a-c and 4a-c**). The overactivation of VCAM-1 and ICAM-1 by LPS in CYB5R3 deficient endothelial cells is through the NF-κB pathway and was completely blunted by simultaneous knockdown of NOX4 (**Figure 2h-i, 3e, and 4d**). A similarly enhanced induction effect by TNF-α was seen for VCAM-1 (**Figure 5a-b**) but not ICAM-1 (**Figure 5c**). A plausible explanation is that temporal regulation of CYB5R3 on TNF-α induced ICAM-1 is different, which was not captured after overnight treatment. More importantly, although both *Vcam-1* and *Icam-1* are NF-κB target genes, it is not a surprise that they can be differentially modulated by other transcription factors and upstream signaling pathways^60, 61^. The exact mechanism defining how CYB5R3 and NOX4 affect LPS and TNF-α induced inflammatory signaling remains a question, and future studies are needed to address specific targeting in the pathway.

As an LPS-induced cytokine, TNF-α contributes to the inflammatory cascade initiated by LPS *in vivo*^40, 41^. To investigate this *in vivo* role of LPS, we created inducible endothelial cell-specific *Cyb5r3* knockout mice to investigate the role of endothelial CYB5R3 in the LPS induced inflammatory cascade (**Figure 1a**). Circulating LPS, as seen in sepsis, activates vascular endothelial cells to express adhesion molecules such as VCAM-1 and ICAM-1, which recruit leukocytes to the vessel. Leukocytes, such as monocytes, express NOS2 that produces nitric oxide and causes extensive vasodilation resulting in systemic hypotension (**Figure 8**). We found that LPS-induced hypotension was exacerbated in mice lacking CYB5R3 in the endothelium (**Figure 1c**). Moreover, without endothelial CYB5R3, aortae isolated from LPS challenged mice showed compromised acetylcholine-induced vasodilation and augmented *Nos2 and Vcam-1* mRNA expression, indicating potentiated inflammatory activation and deteriorated endothelial function (**Figure 1d-f**). Interestingly, endothelial *Cyb5r3* deficiency did not affect LPS induced *Icam-1* transcription, which was similar to the effect of CYB5R3 on TNF-α signaling. Based on our previous work, α-globin is not expressed in large artery endothelium but rather small arteries and arterioles at the myoendothelial junction and modulates endothelial NOS3-derived NO signaling^25^. Therefore, it is unlikely that α- globin contributes to endothelial cell dysfunction in the aorta. However, we cannot rule out the possibility that α-globin may also contribute to the hypotensive effects that may arise from the dilation of smaller resistance arteries. Future studies are needed to elucidate a likely nuanced and relative impact of endothelial CYB5R3 on the LPS signaling cascade in various segments of the vascular tree. We recognize that the increase in O_2_^•-^ by *Cyb5r3* knockdown is inconsistent with its effect in smooth muscle cells^62^. However, the seemingly incongruous results can be caused by different expression levels of NOX isoforms and cellular reliance on mitochondrial respiration for bioenergetics. Therefore, the role of CYB5R3 in redox signaling should be examined tissue by tissue.

The regulation of inflammatory signaling by CYB5R3 can significantly impact the pathological progression of inflammatory disease in patients. CYB5R3 T117S, a mutant form encoded by a polymorphism (CYB5R3 *c.350C>G*), has a high allele frequency (23%) in the African American population^23^. An important question is whether patients carrying CYB5R3 T117S develop more severe symptoms in infectious diseases, such as sepsis, or chronic inflammatory diseases, such as atherosclerosis. It is also worth noting that CYB5R3 is the best-defined reductase among its family members (CYB5R1- 4 and CYB5RL). The endothelial cell expression levels of these proteins vary across tissues according to the Human Protein Atlas single-cell RNA seq database (RNA single cell type tissue cluster data available from https://www.proteinatlas.org). It is not clear whether other reductases may also regulate the NOX family of enzymes in the cardiovascular system.

In conclusion, by carefully examining the subcellular locations of CYB5R3 and NOX4, we determined their subcellular orientation on the mitochondrial outer membrane in primary endothelial cells. Their spatial proximity favors an interaction permitting CYB5R3-derived semiubiquinone radical to facilitate the reduction of O_2_^•-^ generated by NOX4. This novel mechanism contributes to NOX4’s ability to produce H_2_O_2_ rather than O_2_^•-^. Collectively, our current work provides important and novel insight regarding the modulation of NOX4 activity that may have significant translational relevance for the management of inflammatory diseases in patients who are carriers of the CYB5R3 loss-of-function variants.

## ACKNOWLEDGMENT

We thank Simon Watkins at the Center for Biologic Imaging, University of Pittsburgh, for the use of light and electron microscopes.

## SOURCES OF FUNDING

This work was supported by National Institutes of Health (NIH) R01 awards [R01 HL 133864 (A.C.S), R01 HL 128304, R01 HL 149825, R01 HL 153532 (A.C.S), R01 GM 125944 and R01 DK 112854 (F.J.S.)]. American Heart Association (AHA) Established Investigator Award 19EIA34770095 (A.C.S.)], American Heart Association Post-doctoral Fellowship 19POST34410028 (S.Y.), American Society of Hematology (ASH) Minority Hematology Graduate Award (A.M.D-O.) Junta de Andalucía grant BIO-177 (P.N.), the FEDER Funding Program from the European Union and Spanish Ministry of Science, Innovation and Universities grant RED2018-102576-T (P.N.), NIH 1S10OD021540-01 to Center for Biologic Imaging, University of Pittsburgh, NIH 1S10RR019003-01 (Simon Watkins [S.W.]), NIH 1S10RR025488-01 (S.W.), NIH 1S10RR016236-01 (S.W)

## AUTHOR CONTRIBUTIONS

S.Y. and A.S. conceived and designed the study. S.H., S.Y., and A.M.D-O. performed and interpreted animal experiments. S.Y., M.P.M., S.S., and Y.L. performed and interpreted cell culture, molecular biology, and biochemistry experiments. M.J.C., M.S., C.M.S, and D.S. designed, performed, and interpreted super-resolution confocal microscopy and electron microscopy. M.F. and F.J.S. designed, performed, and interpreted O_2_^•-^ measurements. M.R. and S.S. designed, performed, and interpreted experiments on mitochondrial function. P.N. designed, performed, and interpreted CoQ-related experiments. E.C-P. and P.J.P. designed and interpreted NOX4-related experiments. S.Y. K.C.W., F.J.S., P.J.P., and A.C.S. wrote the manuscript, contributed to the writing, or critically reviewed the manuscript. C.M.S., D.S., P.N., S.S., F.J.S., P.J.P., and A.C.S. provided materials for the study. All authors reviewed and approved the manuscript.s

## CONFLICT OF INTEREST

The authors declare that they have no conflict of interest.

## SUPPLEMENTARY METHODS AND MATERIALS

### Telemetric measurements of blood pressure

Mice were anesthetized with inhaled isoflurane before telemetry units (HDX-10, Data Sciences International) were surgically implanted in the left common carotid artery. After recovering for 14 days, systolic blood pressure data were recorded with the manufacturer’s software (Ponemah) for 24 hours to determine the baseline LPS (Escherichia coli O111:B4, Sigma, L2630) was injected intraperitoneally at a sublethal dose (5 mg/kg in saline). To control for the circadian variation of blood pressure, all LPS injections were performed at the same time (2-3 PM), and the recorded systolic pressure was compared to the baseline reading at the same time to determine the change of pressure.

### *Ex vivo* wire myography

After tamoxifen treatments, LPS was administered intraperitoneally at 5 mg/kg. Eight hours after the LPS injection, mice were euthanized, and aorta was collected for myography according to our previous publications^26^. Briefly, thoracic aortae were rapidly excised placed in room temperature physiological salt solution (PSS), cleaned of fat, cut into 2 mm rings, and placed on a two-pin myograph (DMT 620M) filled with PSS containing (mM): NaCl 119, KCl 4.7, MgSO_4_ 1.17, KH_2_PO_4_ 1.18, D-glucose 5.5, NaHCO_3_ 25, EDTA 0.027, CaCl_2_ 2.5, pH 7.4 when bubbled with 95% O_2_ 5% CO_2_ at 37°C. Following a 30-minute rest, aortas were incrementally stretched to 500 mg initial tension. Vessels were then constricted with 60mM KCl for 5 minutes to test the viability and then washed three times with PSS and allowed to rest for 30 minutes. A final wash was performed, and vessels rested for additional 10 minutes.

Following the final 10 min rest period, vessels were constricted with a dose-response of phenylephrine (50uM-1uM). After reaching a plateau, a continuous dose-response curve of acetylcholine (10 nM-100uM) was used to assess endothelium-dependent relaxation. Ca^2+^ free PSS containing 100μM sodium nitroprusside was added to determine maximal dilation.

### Quantification of gene expression

Aorta samples were pulverized in liquid nitrogen with a Kimble Teflone® pestle and motor homogenizer. The pulverized sample was dissolved in Trizol^TM^ reagent (Thermo Fisher Scientific, 15596026). Cultured endothelial cells were directly lysed in the Trizol^TM^ reagent. Total RNA was extracted from Trizol^TM^ with the Direct-zol™ RNA Miniprep Plus kit (Zymo, R2072) according to the manufacturer’s instructions. Reverse transcription was performed using SuperScript™ IV VILO™ Master Mix (Thermo Fisher Scientific, 11756050). Quantitative PCR for the cDNA library (RT-PCR) was set up with Power SYBR™ Green PCR Master Mix (Thermo Fisher Scientific, 4367659) and measured by QuantStudio^TM^ 5 real-time PCR machine (Thermo Fisher Scientific). The delta-delta Ct method was used to analyze mRNA expression levels. Primer sequences used in this study are as the following. Mouse *Actb* (housekeeping gene): forward 5’- CTTTGCAGCTCCTTCGTTG-3’; reverse 5’-CGATGGAGGGGAATACAGC-3’. Mouse *Vcam-1*: forward 5’-GCAAAGGACACTGGAAAAGAG-3’, reverse 5’- TCAAAGGGATACACATTAGGGAC-3’. Mouse *Icam-1*: forward 5’- GCAGAGGACCTTAACAGTCTAC-3’, reverse 5’-TGGGCTTCACACTTCACAG-3’. Mouse *Nos2*: forward 5’-ACATCGACCCGTCCACAGTAT-3’, reverse 5’- CAGAGGGGTAGGCTTGTCTC-3’. Human *Rps13* (housekeeping gene): forward 5’-TCTCCTTTCGTTGCCTGATC-3’, reverse 5’-AATCTGCTCCTTCACGTCG-3’. Human *Vcam-1*: forward 5’-TTCTCATCACGACAGCAACTT-3’, reverse 5’- GGAGTCACCAATCTGAGCAAG-3’. Human *Icam-1*: forward 5’- TGACCGTGAATGTGCTCTC-3’, reverse 5’-CTGTATTTCTTGATCTTCCGCTG-3’. Human *Cyb5r3*: forward 5’-ATGGGGGCCCAGCTCAG-3’, reverse 5’- AGCGCTGGAACAGCTTCAT-3’. Human *Nox4*: forward 5’- TCACAGAAGGTTCCAAGCAG-3’, reverse 5’-ACTGAGAAGTTGAGGGCATTC-3’.

### Immunoblotting

HAECs were lysed directly in 2X Laemmli buffer. HEK293(FT) cells were lysed in radioimmunoprecipitation (RIPA) buffer with proteinase (P8340-5ML) and phosphatase inhibitors (P5726-5ML). Protein samples in RIPA buffer were quantified using Pierce™ BCA Protein Assay Kit (Thermo Fisher Scientific, 23225), diluted, and mixed with an equal amount of 2X Laemmli buffer. All samples were denatured at 95 °C for 10 minutes. Western blotting was performed as we previously described^63^. Proteins were resolved in NuPAGE Bis-Tris gradient gels (Thermo Fisher Scientific, NP0336BOX) and transferred to 0.2 µm nitrocellulose membranes (BIO-RAD 1620112). Proteins of interest were probed with specific antibodies overnight at 4 °C, followed by infrared fluorescent secondary antibodies for one hour at the room temperature. Blots were imaged with LI- COR Odyssey® CLx, and fluorescent intensity was quantified with LI-COR Image Studio. Antibodies and dilutions used in this study are as the following. VCAM-1 (Abcam, ab134047) 1:1,000; ICAM-1 (Santa Cruz, sc-8439) 1:500; α-tubulin (Sigma, T6074) 1:10,000; CYB5R3 (Proteintech, 10894-1-AP) 1:2,500; NF-κB p65 (Cell Signaling, 8242) 1:1,000; Phospho-NF-κB p65 (Ser536) (Cell Signaling, 3033) 1:1,000; NOX4 (Proteintech, 14347-1-AP; or Abcam, ab133303) 1:1,000; TOM20 (Santa Cruz, SC- 11415) 1:2,000; COX IV (Abcam, ab14744) 1:1,000; catalase (Cell Signaling, 12980) 1:1,000; oxidative phosphorylation complex antibody cocktail (Abcam, ab110411) 1:1,000; (SOD1 (Cell Signaling, 2770) 1:1,000; SOD2 (Cell Signaling, ab13533) 1:1,000; IRDye® 800CW Donkey anti-Rabbit IgG Secondary Antibody (LI-COR) 1:15,000; IRDye® 680RD Donkey anti-Mouse IgG Secondary Antibody (LI-COR) 1:15,000.

### Immunofluorescence staining

For tissues, samples were fixed in 4% paraformaldehyde (Santa Cruz, sc-281692) overnight at 4 °C before they were paraffin-embedded and sectioned at a 7-μM thickness. After standard deparaffinization, antigens were retrieved using a heat-mediated citric acid-based method (Vector Laboratories, H-3300). After one-hour blocking with 10% horse serum at the room temperature, sections were incubated with primary antibodies overnight at 4 °C and secondary antibodies for one hour at the room temperature. For cell culture experiments, HAECs were plated on coverslips and allowed to grow confluent. Cells were fixed in 4% paraformaldehyde, permeabilized with 0.1% Triton X-100 (Sigma, X100-500ML), and blocked with 10% horse serum. Subsequently, coverslips were stained with primary and secondary antibodies. Antibodies and concentrations used for immunofluorescent staining are as the following. CYB5R3 (Proteintech, 10894-1-AP) 2 µg/ml; FITC conjugated α-actin (Sigma, F3777) 1:500; NOX4 (Proteintech, 14347-1-AP) 2 µg/ml; TOM20 (Santa Cruz, SC-17764) 2 µg/ml. DAPI (Thermo Fisher Scientific, D3571, 1:100) was used for counterstaining along with the Alexa Fluor secondary antibodies. Fluorescent images were captured using a Nikon A1 confocal microscope or a Leica STED super-resolution confocal microscope at the Center for Biological Imaging at the University of Pittsburgh. For the super-resolution imaging, images were sequentially captured using a 1.4NA 100x oil immersion STED objective with 0.019µm pixels and 0.150 µm z steps. NOX4 and CYB5R3 were excited with a pulsed 555nm laser and depleted with the 775nm STED laser. Emission was captured between 563-651nm and temporally gated between 1- 6nsec. TOM20 was excited with a 663nm laser and depleted with the 775nm STED laser. Emission was captured between 670-750nm and temporally gated between 1-6 nsec.

### In vitro transfection of siRNA and expression vectors

The transfection of siRNA in HAECs and expression vectors in HEK293 cells were achieved using Lipofectamine 3000 (Thermo Fisher Scientific, L3000015) according to the manufacturer’s instructions. Non-targeting siRNA (siNT) and human *Cyb5r3* targeting siRNA (siR3) were purchased from Horizon Discovery (D-001810-01-20 and L-009554-00-0005). Human *Nox4* targeting siRNA was purchased from Invitrogen (Stealth siRNA, HSS121312). The day before transfection, HAECs were seeded at 15,000-20,000 cells/cm^2^ in 12-well plates. For *Cyb5r3* silencing alone, 10 µM of siNT or siR3 was given to HAECs in lipid complex overnight, before the medium was replenished to allow cells to recover. For experiments involving *Cyb5r3* and *Nox4* double knockdown, four groups of cells received 20 µM siNT, 10 µM siNT + 10 µM siR3, 10 µM siNT + 10 µM siNox4, or 10 µM siR3 + 10 µM siNox4, respectively. In this way, siRNA loads were kept the same in all treatment groups.

HEK293(FT) cells were seeded at 80,000 cell/cm^2^ in 12-well plates. Plasmids for NOX4, p22, and CYB5R3 WT were used at 0.3 µg per well. For unknown, CYB5R3 mutants showed different expression efficiency or protein stability. The amount of plasmid used per well was empirically adjusted as the following: K111A 0.15 µg, K111A/G180V 0.2 µg, ΔAA1-23 0.09 µg. When experiments were performed in different cell culture vessels, the plasmid amount is scaled according to the growth area.

Two days after the transfection of siRNA or expression vectors, HAECs were treated with LPS, and HEK293(FT) cells were collected for downstream analysis.

### Lentiviral production and transduction

The shRNA vector against the human *Cyb5r3* 3’-untranslated region (UTR) was purchased from MISSION® shRNA Library (Sigma, TRCN0000236407). For the Lentiviral expression vector for CYB5R3-APEX, the APEX ORF was amplified from pcDNA3-Connexin43-APEX (Addgene, 49385), fused to CYB5R3 in pcDNA3.1, and cloned into pLV-*IRES-GFP*. To generate lentivirus, HEK293FT cells were seeded in 15- cm dishes at 80,000 cells/cm^2^. On the following day, for each plate, 8.4 µg pLP1, 4.8 µg pLP2, 6 µg pLP/VSVG (Thermo Fisher Scientific, K497500), and 15 µg shRNA or expression vector were mixed with 171 µg polyethylenimine. After 20 minutes of incubation, the plasmid mixture was spread on HEK293FT cells and left for 8 hours before the medium was changed. Medium containing lentivirus was collected 2-4 days after transfection, and the virus was precipitated by adding 1/3 the volume of a viral concentrator (40% [w/v] polyethylene glycol 8000 and 1.2 M NaCl in phosphate-buffered saline) and reconstituted in phosphate-buffered saline.

To establish *Cyb5r3* knockdown cell lines, HEK293FT cells were treated with a series of dilutions of lentiviral *Cyb5r3* shRNA. The monoclonal selection was performed with 2 µg/ml puromycin from cells treated with the least effective dose of the virus. CYB5R3 expression was further examined in selected cell lines by immunoblotting. To transiently express CYB5R3-APEX2 in HAECs, cells were treated with diluted expression lentivirus so that 10% of cells were GFP positive.

### Mitochondria isolation

Mitochondria were isolated from cultured cells as previously described^64^. Briefly, a 10- cm dish of HEK293FT cells was dislodged and resuspended in ice-cold hypotonic buffer (10 mM NaCl, 1.5 mM MgCl_2_, 10 mM Tris-HCl, pH 7.5) for 5-10 minutes. Swollen cells were mechanically broken using a Dounce homogenizer with the tight-fitting pestle. Osmolarity was recovered by adding a 2.5X homogenization buffer (525 mM mannitol, 175 mM sucrose, 12.5 mM Tris-HCl, 2.5 mM EDTA, pH 7.5). Nuclei and intact cells were removed by centrifugation at 1,300 g for 5 minutes at 4 °C. Mitochondria were pelleted from the supernatant by centrifugation at 17,000 g for 15 minutes at 4 °C. For immunoblotting, mitochondria were solubilized in RIPA buffer for protein quantification. Subsequently, 20 µg whole cell lysate and 4 µg mitochondrial lysate were loaded on gels.

For proteinase K digestion, two 15-cm dishes of HEK293FT cells were used to get a larger amount of mitochondria. Instead of lysing the pellet, it was gently reconstituted in 1X homogenizer buffer (210 mM mannitol, 70 mM sucrose, 5 mM Tris-HCl, 1 mM EDTA, pH 7.5). Proteinase K was added to aliquots of mitochondria homogenate. After 10 minutes of incubation at room temperature, proteinase inhibitors and Laemmli buffer were added to each aliquot for downstream immunoblotting.

### Measurement of CYB5R3 activity

CYB5R3 activity was measured by the reduction of 2,6-Dichlorophenolindophenol (DCPIP). HEK293FT cells were lysed with proteinase and phosphatase inhibitors in RIPA buffer for protein quantification. All samples were diluted to 200 µg/ml in Tris-HCl buffer (100 mM, pH6.8). In a 96-well plate, 50 µl diluted samples, 100 µl DCPIP (450 µM in 100 mM Tris-HCl, pH 6.8), and 50 µl NADH (1 mM in 100 mM Tris-HCl, pH 6.8) were added for each reaction. Immediately after adding NADH, dynamic absorbance (600 nm) reading was started at 37 °C until reactions plateau. The log phase slope was used to determine the reaction rate.

### Proximal ligation assay

Proximal ligation assay was performed using Duolink In Situ PLA Probe (Sigma, Anti-Rabbit PLUS [DUO92002-30rxn], Anti-Mouse MINUS [DUO92004-30rxn]) according to manufacturer’s instructions with modifications. HAECs were grown on coverslips to confluence. Following methanol fixation, cells were blocked in Duolink blocking buffer for one hour at 37 °C. Primary antibodies, mouse anti-CYB5R3 (Santa Cruz, sc-398043) and rabbit anti-NOX4 (Proteintech, 14347-1-AP), were diluted to 2 µg/ml and added to coverslips for overnight incubation at 4 °C. The next day, coverslips were rinsed and incubated with anti-rabbit plus and anti-mouse minus secondary antibodies for one hour at 37 °C. Ligation and amplification were setup using provided reagents and lasted 30 and 100 minutes, respectively, at 37 °C. Afterward, DAPI was used as the counterstaining. Confocal images were taken with a Nikon A1 microscope at the Center for Biologic Imaging at the University of Pittsburgh and quantified using ImageJ.

### Detailed method on transmission electron microscopy

The specimens were fixed in cold 2.5% glutaraldehyde (25% glutaraldehyde stock EM grade, Taab Chemical, Berks, England) in 0.01 M PBS (sodium chloride, potassium chloride, sodium phosphate dibasic, potassium phosphate monobasic, Fisher, Pittsburgh, PA), pH 7.3. The specimens were rinsed in PBS, post-fixed in 1% osmium tetroxide (Osmium Tetroxide crystals, Electron Microscopy Sciences, Hatfield, PA) with 1% potassium ferricyanide (Potassium Ferricyanide, Fisher, Pittsburgh, PA), dehydrated through a graded series of ethanol (30% - 90% - Reagent Alcohol, Fisher, Pittsburgh, PA, and 100% - Ethanol 200 Proof, Pharmco, Brookefield, CT), and embedded in Poly/Bed® 812 (Luft formulations) (Dodecenyl Succinic Anhydride, Nadic Methyl Anhydride, Poly/Bed 812 Resin and Dimethylaminomethyl, Polysciences, Warrington, PA). Semi-thin (300 nm) sections were cut on a Leica Reichart Ultracut (Leica Microsystems, Buffalo Grove, IL), stained with 0.5% Toluidine Blue in 1% sodium borate (Toluidine Blue O and Sodium Borate, Fisher, Pittsburgh, PA) and examined under the light microscope. Ultrathin sections (65 nm) were examined on JEOL 1400 transmission electron microscope (grant #1S10RR016236-01 NIH for Simon Watkins) with a side mount AMT 2k digital camera (Advanced Microscopy Techniques, Danvers, MA).

### HPLC-MS/MS quantification of 2-hydroxyethidium (2OHE^+^)

The HPLC-MS/MS based measurement was performed according to the previous publication^65^. To prepare the sample, HAECs were transfected in 6-well plates with siRNA for 48 hours. Cells were incubated with 10 μM dihydroethidium (DHE) (Cayman) for 1 hour, washed with ice-old PBS, and scraped in PBS. Three wells of cells were pooled together and pelleted by centrifugation at 1000 g for 5 minutes. The pellet was lysed in PBS containing 0.1% Triton X-100 and 1 μM 3,8-diamino-6- phenylphenanthridine (DAPP) (Sigma). While an aliquot of lysate was used to determine protein concentration, 0.2 M perchloric acid (HClO_4_) was added to the remaining lysate for protein precipitation. After centrifugation (20,000 g, 30 minutes at 4 °C), an equal volume of 1 M phosphate buffer (pH 2.6) was added to the supernatant. For HPLC-MS/MS analysis, 2-hydroxyethidium (2-OH-E^+^) was synthetized as previously reported^66^, and analyzed by HPLC-ESI-MS/MS using an analytical Phenyl-hexyl Luna column (2 × 150 mm, 3 μm, Phenomenex) at a 0.25 ml/min flow rate, with a gradient solvent system consisting of water containing 0.1% formic acid (solvent A) and acetonitrile containing 0.1% formic acid (solvent B). Samples were chromatographically resolved using the following gradient program: 20–50% solvent B (0–7 min); 50-100% in 0.25 min and 1 min at 100% solvent B followed by 4.5 minutes of re-equilibration to initial conditions. Detection of 2-OH-E^+^ was performed using an AB Sciex5000 triple quadrupole mass spectrometer (Applied Biosystems, San Jose, CA) equipped with an ESI probe in positive mode with the following parameters: declustering potential (DP) 60 V, entrance potential (EP) 5 V, collision cell exit potential (CXP) 10 V, and a source temperature of 500 °C. Multiple reaction monitoring (MRM) transitions for 2-OH-E^+^ and its internal standard DAPP were used with the following optimized collision energies (CE): MRM 330.1/300, 286/208 with CE 45 and 40 eV respectively. Results were reported as ratio of analyte peakarea/ I.S. peak area normalized to the protein concentration.

### Seahorse Assay

Cellular oxygen consumption rate was measured by with Seahorse Extracellular Flux (XF96) Analyzer (Seahorse Bioscience, Inc., North Billerica, MA) as previously described. HAECs were transfected with siRNA for 48 hours and then seeded in an Agilent Seahorse XF96 cell culture microplate (Agilent, 101085-004) at 200,000 cells/cm^2^. After attachment, cells were treated with LPS (1 µg/ml) overnight. On the day of experiment, medium was changed to DMEM containing 25 mM glucose, 2 mM glutamine and 1 mM pyruvate. In the Seahorse Analyzer, cells were consecutively treated with oligomycin A (2 µM), trifluoromethoxy carbonylcyanide phenylhydrazone (FCCP) (0.5 µM), antimycin A (Ant A) (2 µM), and rotenone (2 µM). After each treatment, two OCR measurements were recorded. The data was reported as fold changes compared to the baseline (DMEM only) level of control group.

### Mitochondrial membrane potential

HAECs were transfected with siRNA for 48 hours. After overnight LPS (1 µg/ml) treatment, cells were trypsinzed, pelleted, and resuspended in 10 µg/ml JC-1 probe (Thermo, T3168). After 20-minute incubation at 37 °C, cells were subjected to 2-color flow cytometry (514/529 nm and 585/590 nm). The emission ratio of 590/529 was recorded as the measurement of mitochondrial membrane potential.

---

**Supplementary Figure 1.**
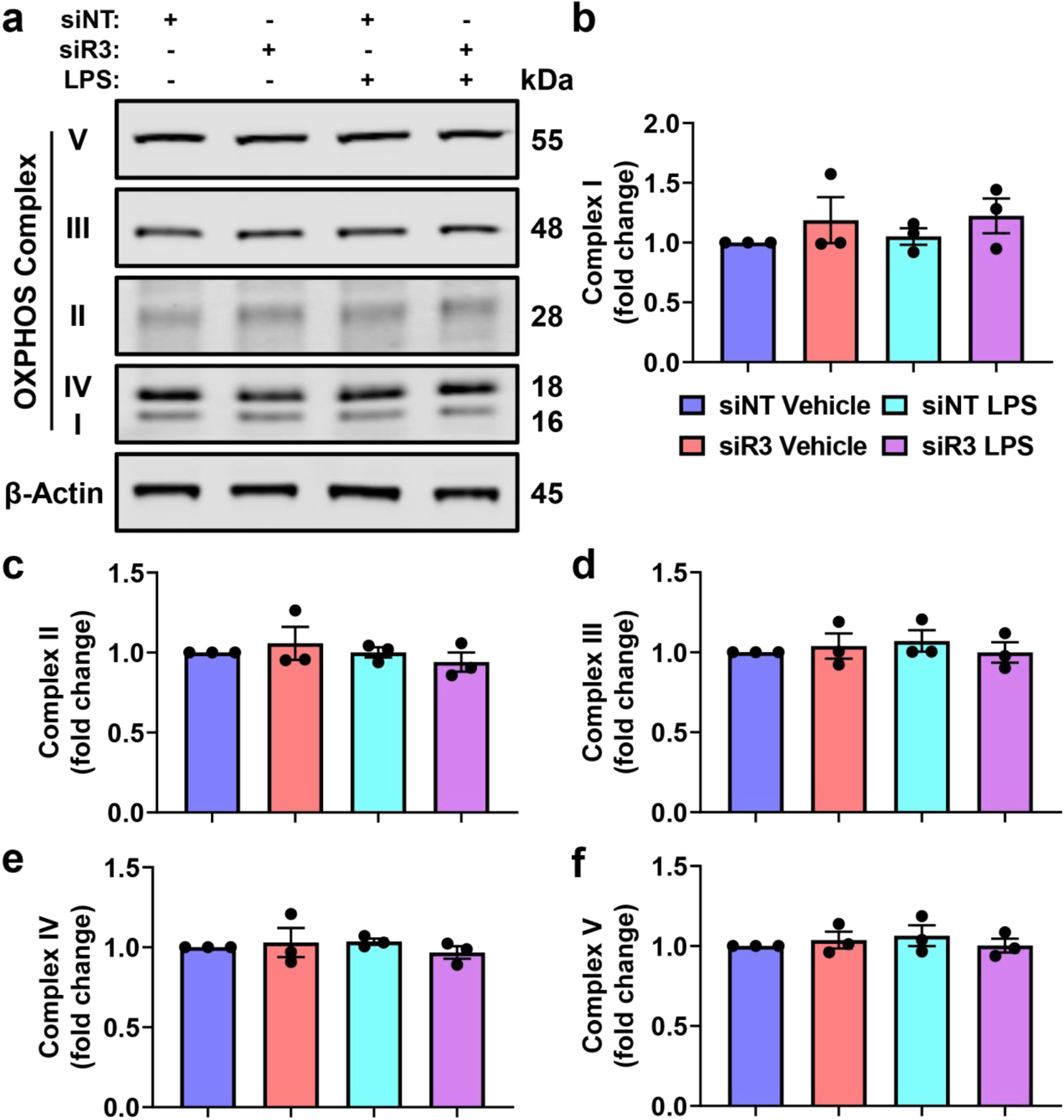
HAECs were transfected with non-targeting (siNT) or *Cyb5r3*-targeting (siR3) siRNA for 48 hours and then treated with LPS (1 µg/ml) overnight. The experiment was repeated 3 times. Representative images were shown in **(a)**, and quantifications of each complex in **(b-f)**. **(b-f)** share the same figure legend.

**Supplementary Figure 2.**
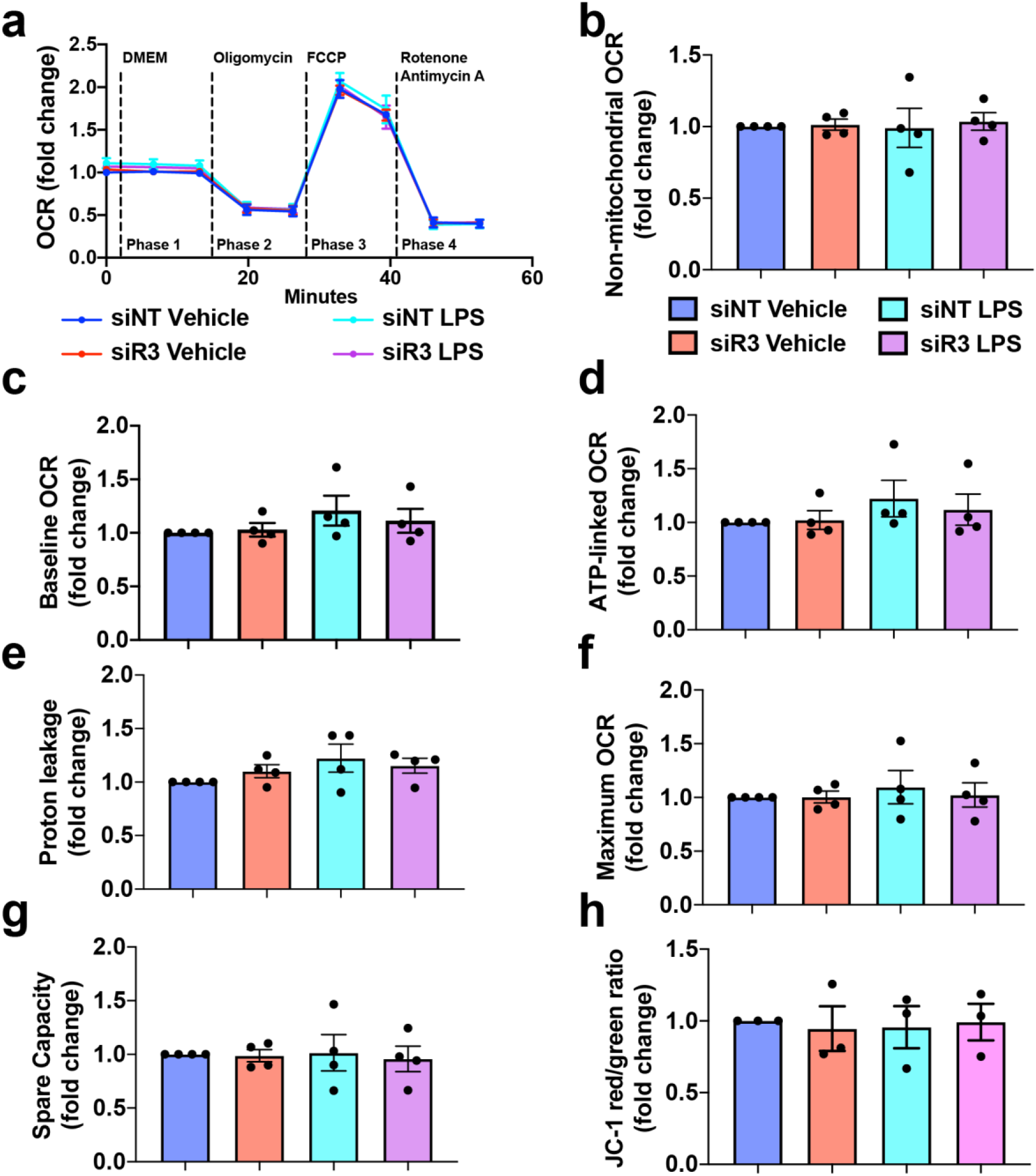
HAECs were transfected with non-targeting (siNT) or Cyb5r3-targeting (siR3) siRNA for 48 hours reseeded in XF96 Seahorse plates, and treated with LPS (1 µg/ml) overnight. The experiment was repeated on different days. **(a)** In each experiment, the oxygen consumption rate (OCR) was normalized to the baseline (Phase 1) level of the control group (siNT vehicle). The curves represent average values from 4 experiments. Each phase corresponds to the OCR under indicated inhibitor treatments for the ease of description. **(b)** Non-mitochondrial OCR = Phase 4. **(c)** Baseline OCR = Phase 1 – Phase 4. **(d)** ATP-linked OCR = Phase 1 – Phase 2. **(e)** Proton leakage = Phase 2 – Phase 4. **(f)** Maximum OCR = Phase 3 – Phase 4. **(g)** Spare capacity = Phase 3 – Phase 1. **(h)** Cells were transfected and treated with LPS in the same way but dislodged for JC-1 probe treatment. The data shows the red (585/590 nm) and green (514/529 nm) ratio averaged from 3 independent experiments. **(b-h)** share the same figure legend.

**Supplementary Figure 3.**
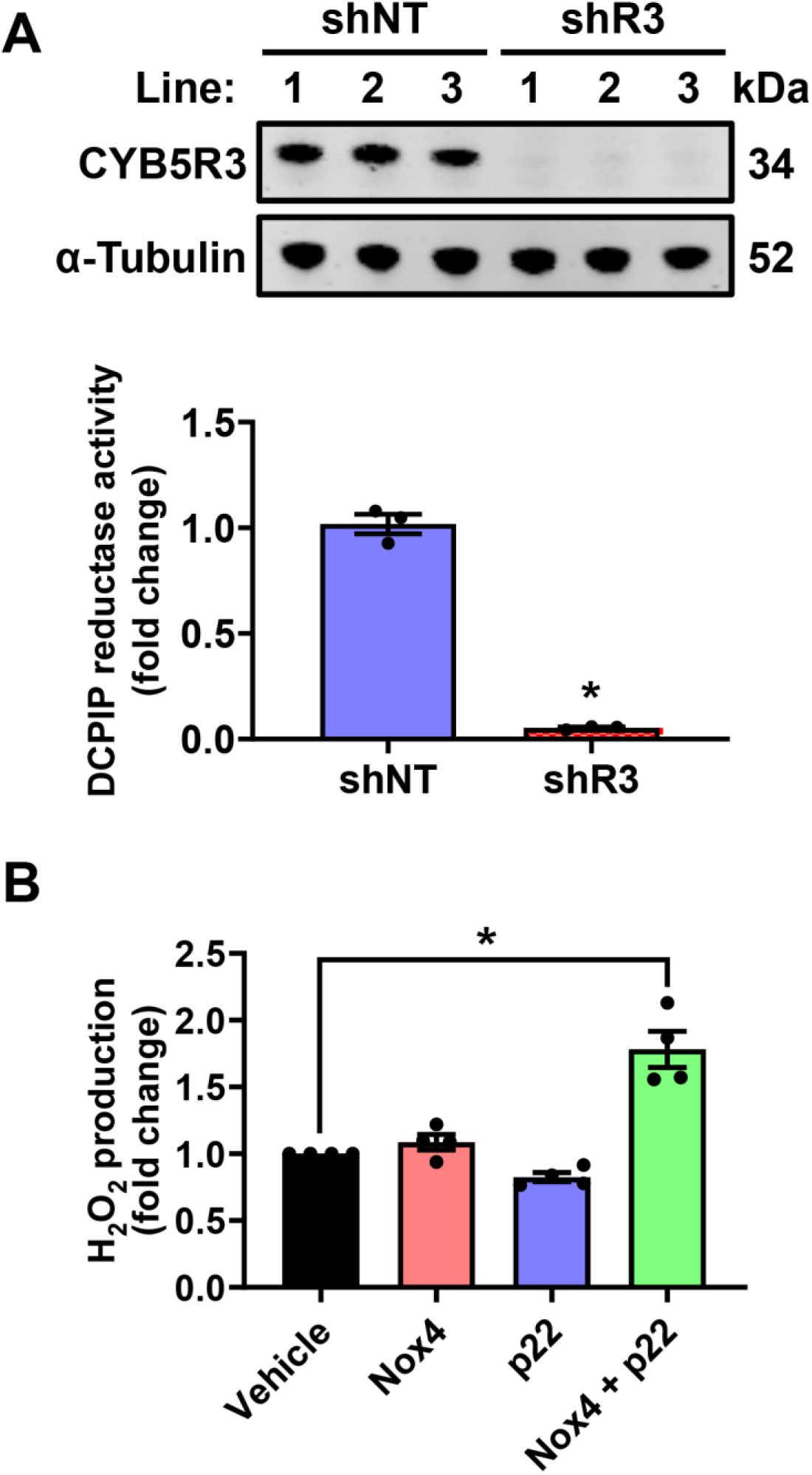
(a) *Cyb5r3* deficient HEK293FT cells were generated using lentiviral shRNA targeting the 3’ untranslated region. After monoclonal selection, three lines of control (non-targeting shRNA treated, shNT) and knockdown (shR3) cells were used to examine CYB5R3 protein expression and activity. N=3 cell lines; * indicates p<0.05 with unpaired t-test. Error bars represent standard deviations. **(b)** HEK293FT cells were transfected with NOX4, p22, or the combination. Only cells expressing both NOX4 and p22 showed elevated H_2_O_2_ production. N=3; * indicates p<0.05 between indicated groups with one-way ANOVA and Dunnett’s multiple comparisons.

**Supplementary Figure 4.**
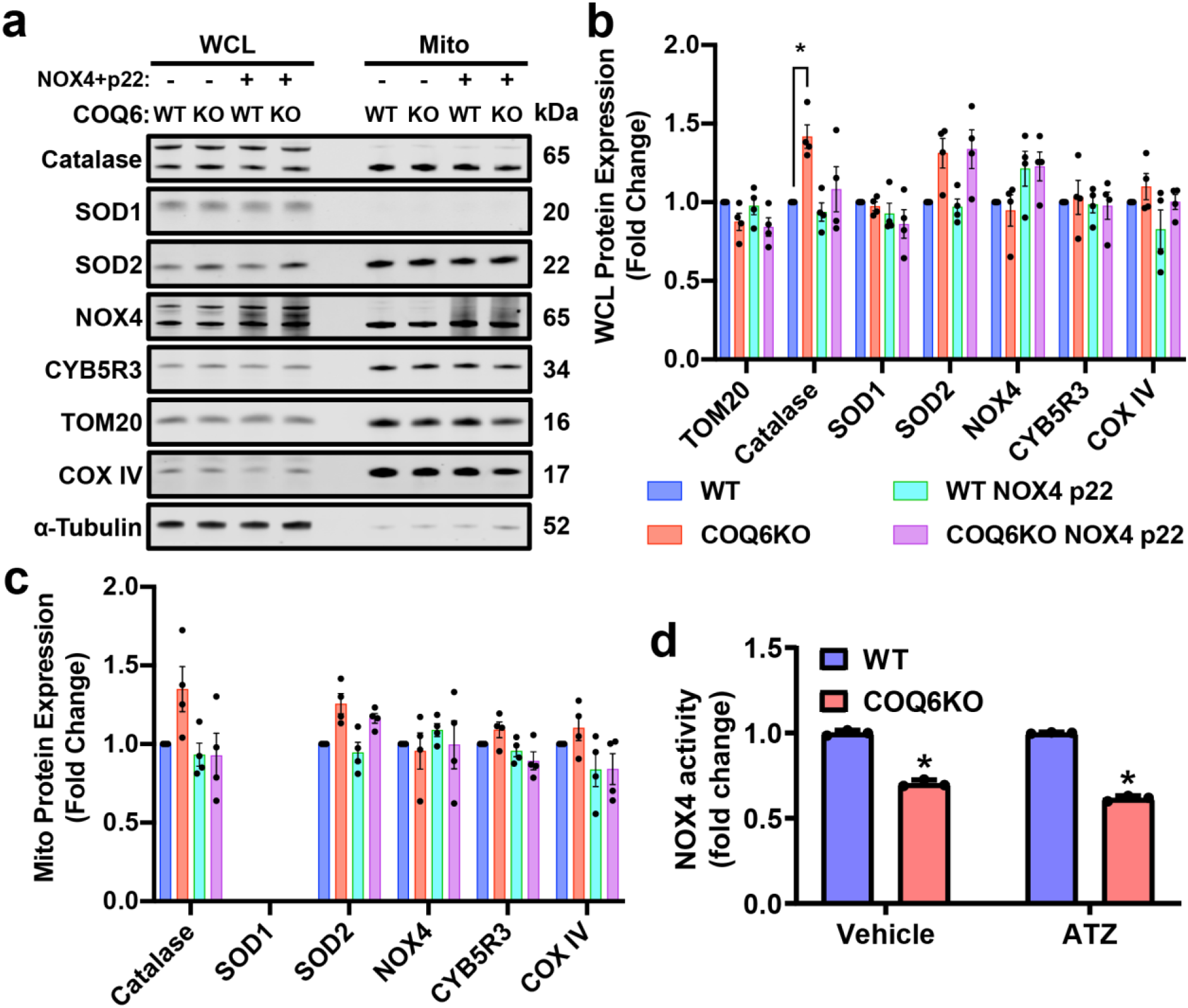
(a) NOX4 and p22 were expressed in wild-type (WT) and *Coq6* knockout (KO) HEK293 cells. The protein expression of catalase, SOD1, and SOD2 were examined in the whole cell lysate (WCL) and isolated mitochondria (Mito). **(b)** The quantification of proteins in WCL normalized to α-tubulin. N=4; * indicates p<0.05 between indicated groups with one-way ANOVA and Tukey’s multiple comparisons. **(c)** The quantification of proteins in mitochondria normalized to TOM20. N=4. **(d)** Wildtype (WT) and *Coq6* knockout (COQ6KO) HEK293 cells were treated with 20 mM 3-amino-1,2,4-triazole (ATZ) immediately prior to H_2_O_2_ measurement. The catalase inhibitor did not increase NOX4-derived H_2_O_2_ in COQ6KO cells. N=3 replicates; * indicates p<0.05 with unpaired t-test; error bars represent standard deviations.

**Supplementary Table 1.**
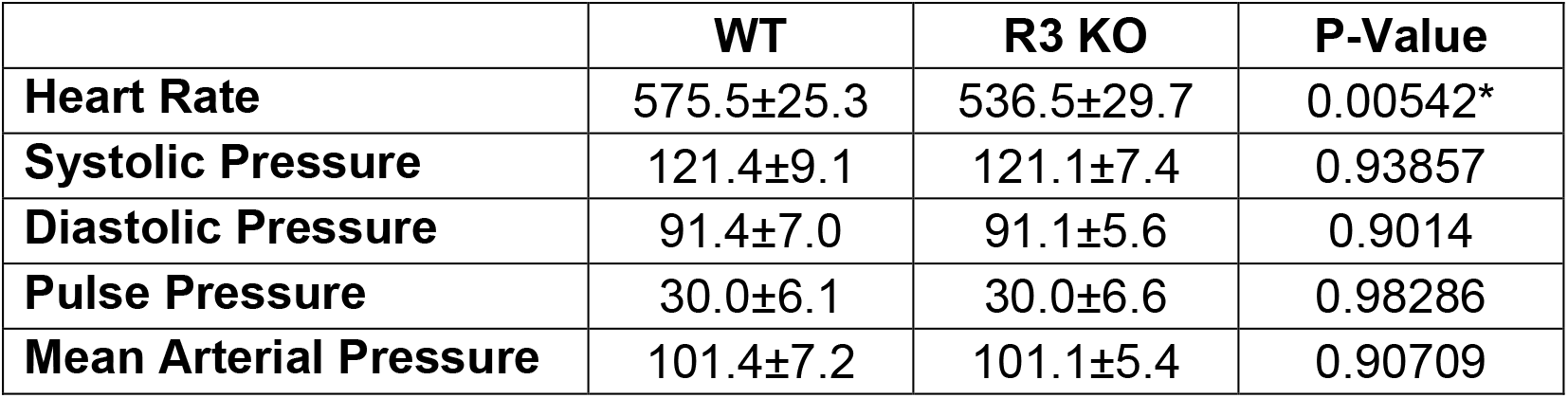
Baseline parameters measured by radio telemetry. Multiple parameters were monitored and recorded by radiotelemetry within the same time-frame, as shown in Figure 1B, and presented as the 24-hour average ± standard deviation. * indicates p<0.05 between tamoxifen-treated wild-type (WT, n=8) and inducible endothelial *Cyb5r3* mice (R3 KO, n=14).

|  | **WT** | **R3 KO** | **P-Value** |
| --- | --- | --- | --- |
| **Heart Rate** | 575.5±25.3 | 536.5±29.7 | 0.00542* |
| **Systolic Pressure** | 121.4±9.1 | 121.1±7.4 | 0.93857 |
| **Diastolic Pressure** | 91.4±7.0 | 91.1±5.6 | 0.9014 |
| **Pulse Pressure** | 30.0±6.1 | 30.0±6.6 | 0.98286 |
| **Mean Arterial Pressure** | 101.4±7.2 | 101.1±5.4 | 0.90709 |

